# Variation in behaviour of native prey mediates the impact of an invasive species on plankton communities

**DOI:** 10.1101/2022.06.02.494478

**Authors:** Sarah S. Hasnain, Shelley E Arnott

**Affiliations:** Department of Integrated Sciences and Mathematics, Habib University, Block 18, Gulistan-e-Jauhar University Avenue, Off Shahrah-e-Faisal, Karachi – 75290, Sindh, Pakistan; Biology Department, Biosciences Complex, Queen’s University, 116 Barrie Street Kingston, ON, Canada K7L 3J9

**Author notes:** **Author contributions** SSH and SEA developed the original idea and experimental system; SSH and SEA contributed to experimental design; SSH carried out the experiment; SSH ran all analyses; SSH wrote the first draft and both authors revised it.

**Keywords:** Intraspecific trait variation, Anti-predator behaviour, *Daphnia* vertical position, Trophic cascade, *Bythotrephes cederströmii*

## Abstract

Population-level differences in predator trait expression influence predator impacts on prey species, altering ecological interactions and trophic dynamics. However, the effect of inter-population differences in prey traits on the impacts of predation on ecological communities remains poorly understood, especially for introduced predators where differences in prey traits could influence the outcome of biological invasions. We examined if differences in *Daphnia* vertical position influenced the impacts of the invasive predator *Bythotrephes cederströmii* on major zooplankton and algal groups. Our results show differences in *Daphnia* vertical position influenced *Bythotrephes* impacts on small cladocerans. Larger reductions in density were observed in mesocosms with greater proportion of hypolimnetic *Daphnia*. Larger increases in algal biomass were also observed in invaded mesocosms with greater proportion of hypolimnetic *Daphnia*. These results suggest that differences in *Daphnia* vertical position influence the magnitude and type of *Bythotrephes* impacts on zooplankton communities.

## Introduction

Predation is an important mechanism structuring aquatic food webs (Paine, 1966; Polis and Holt, 1992). Predators impact prey population dynamics by directly reducing prey densities and imposing strong selective pressure on prey trait expression, often inducing behavioural, morphological and life history changes (reviewed in Lima, 1998; Tollrian &Harvell,1999; Benard, 2004), leading to indirect effects on food web structure and ecosystem function (Paine, 1980; Carpenter et al., 1985; Schmitz et al., 2004; Trussell et al., 2003). For example, reductions in prey density due to predation or change in prey trait expression can increase resource availability, leading to trophic cascades (Paine, 1980; Carpenter et al., 1985; Trussell et al., 2003; Schmitz et al., 2004). However, this research assumes that mean trait values or species identity sufficiently characterize predator-prey interactions, ignoring potential effects of intraspecific variation in both predator and prey traits on ecological communities.

Within a species, individuals can vary across a variety of behavioral, morphological, physiological and life history traits. For species occupying habitat along wide environmental gradients or with large spatial and temporal heterogeneity, variation in selection pressures can result in differences in trait expression, known as phenotypic divergence (reviewed by Crispo, 2008; Pfennig et al., 2010). From an evolutionary perspective, these trait differences are a first step in ecological speciation (reviewed by Dieckmann et al., 2004; Pfennig et al., 2010). Considerably less attention has been paid to the ecological consequences of trait differences between populations. Studies show that population-level differences can influence species abundance, community structure (Post et al., 2008; Palkovacs and Post, 2009; Ibarra-Isassi et al., 2022) and the type and magnitude of food web interactions (Post et al., 2008; Benesh and Kalbe, 2016; Salo et al., 2020). For example, residence duration and foraging morphology differences between two alewife, *Alosa pseudoharengus,* populations impacted prey biomass and the strength of trophic cascades (Post et al., 2008). Most studies examine ecological impacts of inter-population trait variation through predator-prey interactions (but see Benesh and Kalbe, 2016), focusing on the consequences of differences in predator traits. While many laboratory studies have shown that prey trait variation impacts predator-prey dynamics, these effects have rarely been assessed in a community context (but see Lenhart et al., 2018).

The introduction of predators outside their native range results in greater negative impacts on prey communities as compared to native predators (Elton, 1958; Mack et al., 2000; Sih et al., 2010) as prey cannot detect or respond appropriately to the predator due to the lack of shared evolutionary history with the introduced predators (Cox and Lima, 2006; Banks and Dickman, 2007; Sih et al., 2010; Carthey and Banks, 2014; Carthey et al., 2017). These studies assume that introduced predator impacts on native prey are uniform across the invaded range, despite the ubiquity of intraspecific and inter-population variation in traits across ecological systems (Bolnick et al., 2011). Prey naiveté to introduced predators depends on many factors, including the ecological novelty of the predator, the suite of anti-predator defenses available, and the degree of specialization of predator recognition templates and anti-predator defenses possible (Carthey and Banks 2014). Furthermore, inter-population differences in behavioural, morphological, and life history traits may confer protection to some prey, but not others. In lakes invaded by the spiny water flea, *Bythotrephes cederströmii* (formerly *longimanus* (Korovchinsky and Arnott, 2019; hereafter *Bythotrephes*), an Eurasian visual zooplankton predator (Pangle and Peacor, 2009; Jokela et al., 2013) spatially restricted to the upper light penetrating regions of the water column (Pangle and Peacor 2006; Pangle et al., 2007), predation is only expected to directly impact prey populations spatially overlapping with this predator.

*Bythotrephes* is a voracious zooplanktivore (Lehman et al., 1997). Its introduction into North American freshwater systems has resulted in devastating impacts on native zooplankton communities, reducing biomass and species diversity, especially for cladocerans (Yan et al., 2001; Boudreau and Yan, 2004; Barbiero and Tuchman, 2004, Strecker and Arnott, 2005; Strecker et al., 2006; Kelly et al., 2012; reviewed by Azan et al., 2015; Kerfoot et al., 2016). Strong predation on cladocerans, particularly *Daphnia* which are important grazers (Vanderploeg et al., 1993; Schulz and Yuritsa, 1999), has been linked to trophic cascades leading to increased phytoplankton biomass in invaded lakes (Strecker and Arnott, 2008; Walsh et al., 2016). In some invaded lakes, *Daphnia* (Pangle &Peacor 2006; Pangle et al., 2007; Jokela et al. 2011; Bourdeau et al., 2013; Hasnain and Arnott 2019) reside in darker portions of the water column, where *Bythotrephes* predation is reduced (Pangle and Peacor 2009; Jokela et al., 2013). This deep vertical position allows these *Daphnia* to avoid *Bythotrephes* predation, with potential consequences for food web functioning. Despite a large degree of variation in daytime vertical position observed for *Daphnia* populations in natural systems (De Meester, 1993; Tessier and Leibold, 1997; Boeing et al., 2006), the influence of differences in *Daphnia* vertical position on *Bythotrephes* impacts in freshwater ecosystems remains unknown.

Our goal was to determine if differences in *Daphnia* vertical position influence the effect of *Bythotrephes* on plankton communities in invaded lakes. To accomplish this, we manipulated *Bythotrephes* presence in mesocosms with three-tier food webs which were stocked from lakes where *Daphnia* populations exhibited different mean vertical positions and examined if the impacts of *Bythotrephes* on *Daphnia*, other cladocerans, copepod zooplankton and algae abundance differed across mesocosms with different daytime vertical distributions. We expected *Bythotrephes* predation on *Daphnia* to be greater in mesocosms with greater spatial overlap with *Bythotrephes,* i.e. shallow vertical position as compared to those with reduced spatial overlap, i.e. deep vertical position. Greater *Bythotrephes* predation on other cladoceran species was expected in mesocosms with less spatial overlap between *Daphnia* and *Bythotrephes*, resulting in reduced abundances of these species. We expected increases in calanoid copepod abundance in invaded mesocosms regardless of *Daphnia* vertical position due to reduced grazing competition as a result of *Bythotrephes* predation on *Daphnia* and other cladoceran grazers. We predicted that *Bythotrephes* predation on these cladoceran grazers combined with reduced *Daphnia* grazing in mesocosms with deep *Daphnia* vertical position would result in a trophic cascade leading to increased algal biomass.

## Methods

### Study Site and Experimental Design

From July 7^th^ to August 1^st^ 2014, we conducted a field mesocosm experiment to assess the influence of differences in *Daphnia* vertical position on effects of *Bythotrephes* predation on plankton communities. *Bythotrephes* impacts on zooplankton community structure and algal production can occur over short periods of time comparable to the length of our experiment (Strecker and Arnott, 2005; Jokela et al., 2017; Arnott and Azan 2017). Mesocosms were set up in Fletcher Lake (45.20.452’N, 78.47.798’W) Haliburton County, Ontario Canada (Table 1), where *Bythotrephes* was first detected in 2006 (Cairns, 2007). Five mil food grade polythene enclosures, 1m in diameter, 13 m deep (Filmtech Plastics, Brampton, Ontario) and filled with 10990L of water, were closed at the bottom and suspended from floating wooden frames anchored in the lake. Each mesocosm was filled with water pumped from 1.5m and filtered through an 80-µm mesh to exclude crustacean zooplankton, but allow most phytoplankton and some small rotifers and nauplii to pass through. Each mesocosm was covered with screen mesh to prevent colonization by aerial insects.

**Table 1:**
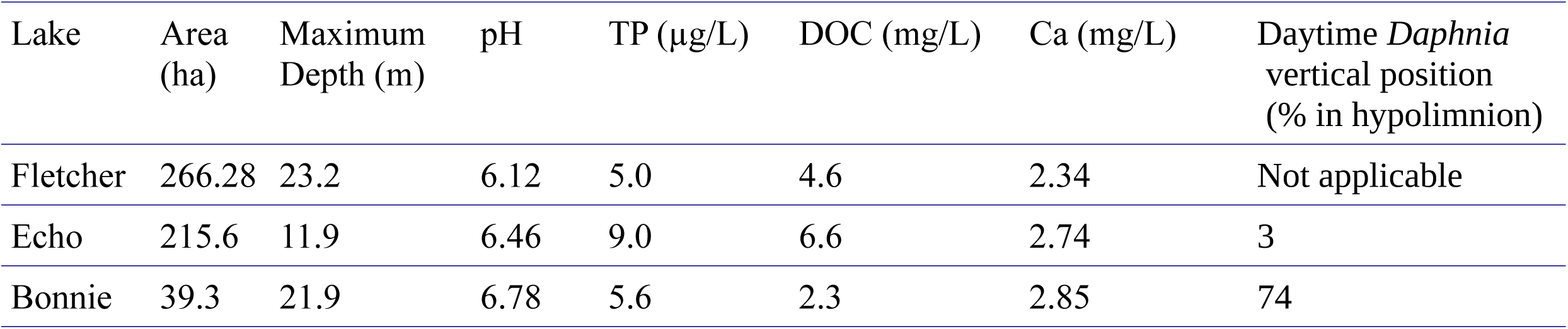
Area, maximum depth, pH, average total phosphorus (TP), dissolved organic carbon (DOC) and calcium (Ca) for Fletcher, Echo and Bonnie lakes based on data from the Canadian Aquatic Invasive Species Network (CAISN) surveys (2008, 2011). Mesocosms were suspended in Fletcher Lake. Zooplankton were stocked from either Echo, Bonnie or combination of both lakes. *Daphnia* daytime vertical position (%) is based on data collected by Hasnain and Arnott (2019). Weekly temperature measurements for mesocosms are provided in Table S1.

To assess if differences in *Daphnia* vertical position influences *Bythotrephes* predation on plankton communities, we manipulated *Bythotrephes* presence and absence in mesocosms across a gradient of *Daphnia* vertical position. We used thermal gradients in lieu of light measurements to determine *Daphnia* vertical position in our mesocosms. With 4.6mg/L of dissolved organic carbon (DOC) present in Fletcher Lake (CAISN 2011), only 1% of ultra-violet and photosynthetically active radiation (PAR) was estimated to be present at 4m (based on average summer solar radiation for south-central Ontario (NASA Atmospheric Science Data Center), and relationships between DOC and light attenuation (Williamson, 2009)). Therefore, we expected no light to be present in the hypolimnion (9-13m) and quantified *Daphnia* vertical position based on their numeric proportion in this region.

*Bythotrephes* predation was expected to be completely absent in the hypolimnion, where light penetration does not occur, as it is a visual predator requiring light to feed (Jokela et al., 2009). After allowing phytoplankton assemblages to increase in biomass without the presence of large zooplankton (>80µm) for three days, mesocosms were stocked with the entire zooplankton community from two uninvaded lakes; Bonnie Lake (45.17.36’N 79.06.45’W; Bracebridge Municipality, Table 1) and Echo Lake (45.17.36’N, 79.06.45’W; Lake of Bays Municipality, Table 1), in south-central Ontario that exhibited contrasting daytime *Daphnia* vertical position. Each mesocosm was randomly inoculated with an ambient density of zooplankton sampled from the same volume of water from either Echo or Bonnie lake, or half volumes for mesocosms stocked from both lakes, resulting in 16 mesocosms stocked from Echo Lake, 16 mesocosms stocked from Bonnie Lake and eight mesocosms stocked from both lakes. This study was part of a larger experiment with additional treatment was applied to Echo and Bonnie mesocosms three weeks after the start of the experiment. Any data collected after the third week was not included in this study.

Half of the mesocosms were randomly assigned the invasion treatment. *Bythotrephes* individuals were collected from Fletcher Lake and Lake of Bays, Muskoka, Ontario (45°15.00’N, 79°04.00’W) using an 80 µm mesh net, and stocked at a density of 10 individuals per m^3^ of epilimnion volume (23 individuals per mesocosm) at the beginning of the experiment. Zooplankton communities in all mesocosms were acclimated for one week prior to *Bythotrephes* addition. There were no *a priori* differences in the proportion of hypo- and non-epilimnetic (hypo- and metalimnetic) *Daphnia* between invaded and uninvaded mesocosms in week 0 (Figure S2, Gamma GLMs, hypo: p = 0.911; non-epi: p = 0.792). *Daphnia* daytime vertical position remained stable across mesocosms throughout the duration of the experiment (see supplementary materials for analysis details, Figure S4).

There were also significant differences in the abundance of zooplankton between Echo and Bonnie lakes, which influenced initial zooplankton densities in our mesocosms. Overall zooplankton density was greater in enclosures stocked from Echo Lake as compared to Bonnie Lake (Figure S1); log normally distributed linear model with lake origin (either Echo, Bonnie, or Both) as a predictor variable and zooplankton density as a response variable (p <0.0001, Echo: 2.490±1.040, Bonnie: 1.497±0.974, Both: 2.910±0.861, see Table S2 for initial densities in each mesocosm). There was no difference in zooplankton starting densities between mesocosms assigned *Bythotrephes* presence and absence treatments (Figure S5; Absent:1.954±1.026, Present: 2.287±1.126 linear model, p>0.05 for all major taxonomic groups). There was very little variation in total algal densities between the mesocosms at the start of the experiment (mean = 6.438±0.110 µg/L, Table S2).

### Sampling protocol and identification

All zooplankton samples were collected between 10am and 3pm. Each mesocosm was sampled prior to the addition of *Bythotrephes* (week 0) to determine abundance and depth distribution of zooplankton and phytoplankton. Samples were also taken at the end of the study (week 3). Epi-, meta- and hypolimnetic boundaries were determined weekly using the thermal profile of Fletcher Lake using a YSI model 600 OMS V2. Zooplankton samples from each mesocosm were collected by towing a closing net with an 80 µm mesh (15 cm diameter) through each thermal layer (starting 20cm above the enclosure bottom for the hypolimnion). Samples were preserved in 90% ethanol. Water samples were collected from the middle of the meta- and hypolimnion using a 2L Van Dorn sampler in weeks 0 and 3. For the epilimnion, water samples were collected from 10 cm below the surface of the mesocosm by submerging the sample container. We determined total algal biomass as well as biomass of green algae, cyanobacteria, diatoms and cryptophytes spectrophotometrically by analyzing a well-mixed 25ml subsample within 24 hours of sample collection (BBE moldaenke Algae Lab Analyser; Schwentinental, Germany).

Zooplankton were enumerated using sub-samples of a known volume and identifying all individuals within each subsample until no new species were found three sub-samples in a row. A minimum of seven sub-samples was counted for each thermal layer in each mesocosm. All specimens were identified to the species level (Ward and Whipple, 1959; Smith and Fernando, 1978; De Melo and Hebert, 1994; Witty, 2004; and Haney et al., 2013). We grouped *Bosmina freyi* and *Bosmina liederi a*s “*Bosmina freyi/liederi”* and *Daphnia pulex a*nd *Daphnia pulicaria a*s “*Daphnia pulex/pulicaria”* due to morphological similarities between these species. For *Daphnia* species, only adults were enumerated. Juvenile copepods were identified as either nauplii or copepodites, without distinguishing between cyclopoid and calanoid juveniles as both have similar diets and occupy a similar trophic position (Finlay and Roff, 2004).

### Statistical analysis

All analyses were conducted in R v 3.2.4 (R Core Development Team 2016) using bbmle v 1.0.17, glmmADMB v 0.7.7, fitdistrplus 1.0.7, piecewiseSEM v 2.1.2, robustbase v 0.93-3 with α = 0.05. Because of differences in starting densities between mesocosms stocked from Bonnie, Echo or both lakes (Table S2, Figure S1), we standardized the change in density between weeks 0 and 3 by calculating per capita change in density for each species and functional group per mesocosm; calculated as density in week 3 divided by density in week 0, which may be sensitive to sampling and demographic effects (see supplementary information). It is possible that the proportion of total hypolimnetic *Daphnia* may have changed during the experiment due to differences in abiotic and biotic variables between the lakes of origin and Fletcher Lake. We found no significant differences in the proportion of hypolimnetic *Daphnia* between the beginning (week 0) and the end of the experiment (week 3, see supplementary information for statistical assessment details). We separately assessed *Bythotrephes* impacts on *Daphnia,* small (<0.85mm) and large cladocerans (>1.0mm) excluding *Daphnia* as well as calanoid, cyclopoid, and juvenile copepods (see supplementary materials for details on zooplankton categorization).

We calculated the density of each species in each thermal layer for every mesocosm by estimating the total number of individuals present in sub samples. Species-specific density in each layer was calculated by dividing the total number of individuals by the volume sampled with a single vertical tow. To calculate the total density of a species in a mesocosm, we summed the total number of individuals in the epi-meta- and hypolimnion samples for each mesocosm and divided this by the total volume of the mesocosm that would be sampled in a single vertical tow. We could not conclusively state whether the absence of a species in our sample represented a true absence in our mesocosm as we could only sample 2% of our mesocosm volume (230L) for zooplankton and < 1% for phytoplankton. We addressed this by adding minimum detection densities to all taxa assessed (see supplementary information).

We used piecewise structural equation modelling (piecewise SEM, Shipley, 2000; Grace 2006) to explore causal relationships between per capita change in density of major zooplankton groups (see above), total algae biomass in week 3, *Bythotrephes* presence, and the proportion of total hypolimnetic *Daphnia* in week 0 (see supplementary information for SEM implementation details). Paths with significant causal relationships (p <0.05) between per capita change in density a zooplankton group or final algal biomass, *Bythotrephes* presence, and *Daphnia* vertical position were re-fit with either normal or gamma distributed generalized linear models to assess the possibility of a significant interaction between the explanatory/exogenous variables. Piecewise SEMs were also fit for the most common species in each zooplankton group with statistically significant causal relationships detected in the full model, with *Bythotrephes* presence, and proportion of total hypolimnetic *Daphnia* in week 0 as exogenous variables, excepting paths for the most common calanoid and cyclopoid species where per capita change in copepodite density was also included. Significant causal relationships in piecewise SEMs for both common species were also re-fit with GLMs to assess if there was a significant interaction between *Daphnia* vertical position and *Bythotrephes* presence. We also refit piecewise SEMs with *Bythotrephes* presence and the proportion of hypolimnetic individuals for *Daphnia* species whose per capita change in density was identified as significantly impacted by *Bythotrephes* presence in earlier SEMs to assess if any effect of proportion of total hypolimnetic *Daphnia* observed was driven by these species. Separate piecewise SEMs were also fit for week 3 biomass of the four major algal groups; green algae, cyanobacteria, diatoms, and cryptophyta using the protocol outlined for the full model, with the nested version also confirmed using AIC.

For all GLMs, we used AICc to assess fit between gamma and log-normal distributions. Model fit was also assessed by visually examining plots of residual versus fitted values and square root of the standard deviance of residuals versus fitted values. Cook’s distance was used to identify influential points (leverage > 1.0). Minimum adequate models were chosen using log-likelihood ratio tests based on Crawley’s (2005) procedure. If influential points were detected, gamma or log-normal robust GLMs were fitted using Mallows or Huber type robust estimators (Cantoni and Ronchetti, 2001; Cantoni and Ronchetti, 2006), which down weights the effect of influential points on model fit. For robust GLMs, minimum adequate models were chosen using Wald-type tests. We did not find any effect of density dependent effects of *Bythotrephes* predation due to differences in starting densities (details about statistical methods used provided in supplementary materials).

## Results

Twenty-eight zooplankton species were present, including six *Daphnia* species (*D. pulex/pulicaria*, *D. ambigua*, *D. catawba*. *D. mendotae, D. parvula,* and *D. dubia*) during our experiment. The most common species (>70% presence across all mesocosms) were *D. catawba* and *D. mendotae* for *Daphnia*, *B. freyi/leideri, E. tubicen,* and *E. longispina* for small cladocerans, *H. glacialis* for large cladocerans, S*kistodiapotmus oregonensis* for calanoids, and *Cyclops scutifer* for cyclopoids. Piecewise SEMs showed that per capita change in density of *Daphnia (Bythotrephes* standardized estimate (SE): -0.11, Vertical position SE: 0.06), small cladocerans (*Bythotrephes* SE: - 0.01, Vertical position SE: 0.008), and cyclopoids (*Bythotrephes* SE: -0.11, Vertical position SE: -0.28) was predicted by *Bythotrephes* presence and the proportion of total hypolimnetic *Daphnia* in week 0 (Table S5, only significant paths presented in Figure 2). Per capita change in juvenile copepod density was only predicted by proportion of total hypolimnetic *Daphnia* (Vertical position SE: 0.21, Figure 2a, Table S5). Change in total algal biomass was predicted by per capita change in density of large cladocerans and juvenile copepods, and the proportion of hypolimnetic *Daphnia* (Vertical position SE: 0.05, Large cladocerans SE: 0.06, Juvenile copepods SE: -0.05, Figure 2a, Table S5).

**Figure 1:**
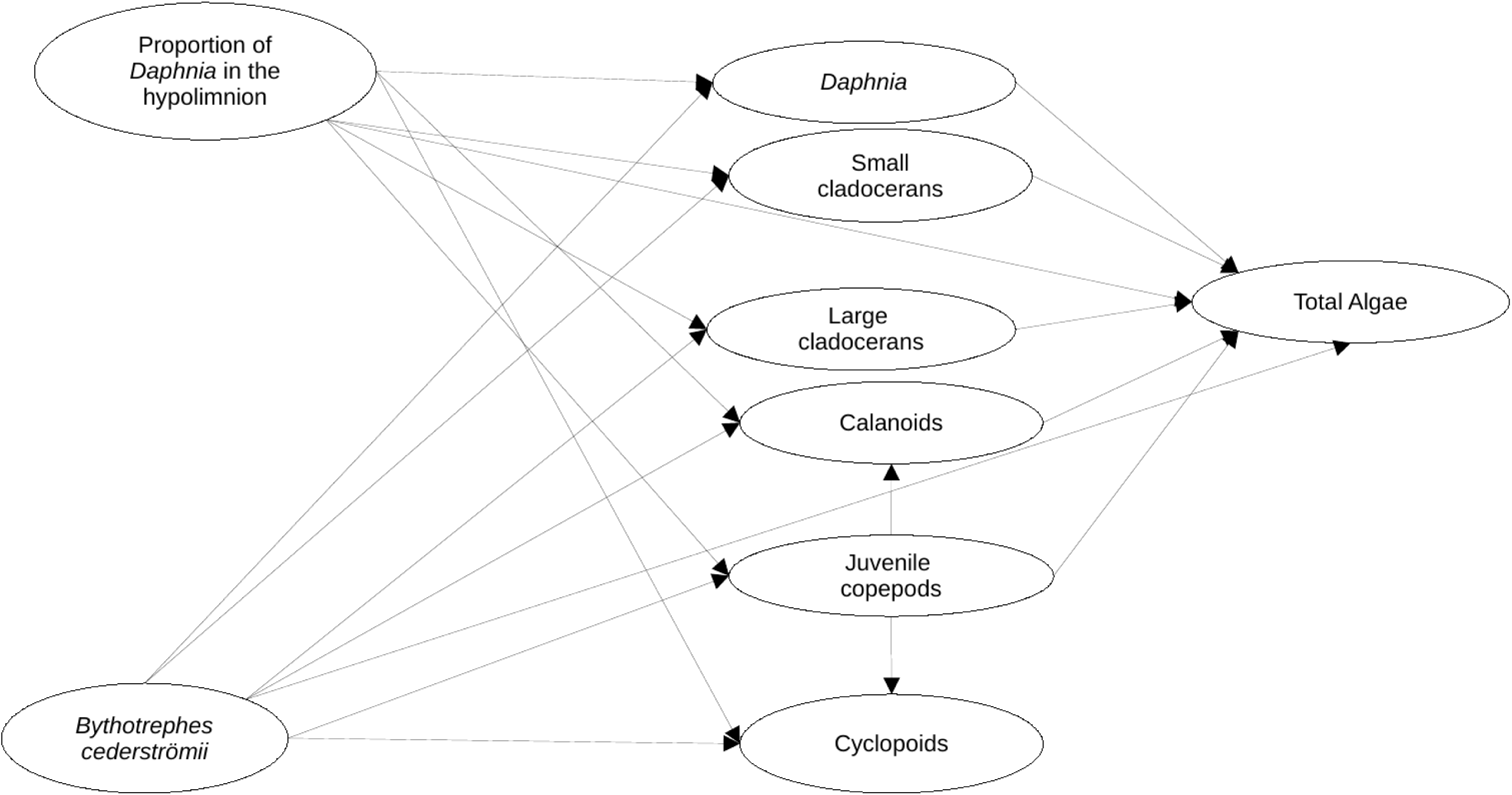
Visualization of the structural equation model (full model) used to assess the impacts of proportion of hypolimnetic *Daphnia* and *Bythotrephes cederströmii* presence on per capita change in *Daphnia*, small and large cladocerans, calanoids, cyclopoids, and juvenile copepods. Model assumes that all paths (represented by arrows) between per capita change density of each zooplankton group, the proportion of hypolimnetic *Daphnia* and *Bythotrephes* presence are possible. For total algal biomass, the path includes all zooplankton groups except cyclopoids, *Bythotrephes* presence, and proportion of hypolimnetic *Daphnia*.

**Figure 2:**
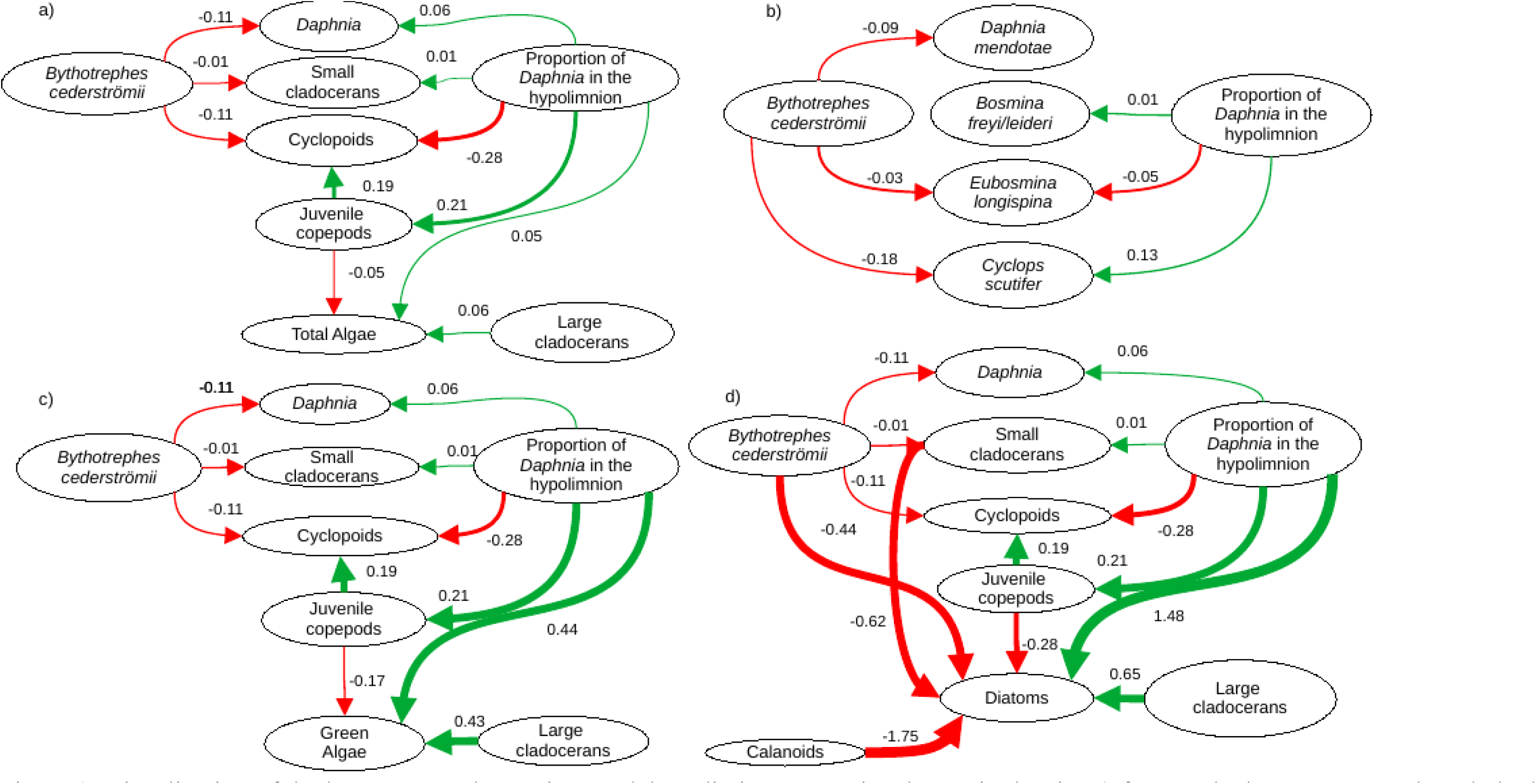
Visualization of the best structural equation model predicting per capita change in density a) for zooplankton groups and total algal biomass in week 3, b) most common *Daphnia*, small cladoceran and cyclopoid species, c) green algal biomass in week 3, and d) diatom biomass in week 3. Arrows represent standardized path coefficients that are statistically significant (p <0.05) with standardized coefficient value. Arrow width is scaled with the size of the coefficient. Arrow colour signifies positive (green) and negative (red) path coefficients.

The proportion of hypolimnetic total *Daphnia* in both invaded and uninvaded mesocosms did not change during our experiment (Uninvaded Week 0 – Week 3: p = 0.07, Invaded Week 0-Week 3: p = 0.09, df = 76, Table S14). The effect of *Bythotrephes* on per capita change in total *Daphnia* density was influenced by the proportion of total hypolimnetic *Daphnia* in week 0 (Gamma GLM, Figure 3a, p = 0.02, df = 36, Table S13). In uninvaded mesocosms with a greater proportion of hypolimnetic *Daphnia*, per capita increase in total *Daphnia* density was larger as compared to mesocosms with fewer hypolimnetic *Daphnia*. Invaded mesocosms had a smaller per capita increase compared to uninvaded mesocosms (Figure 3a).

**Figure 3:**
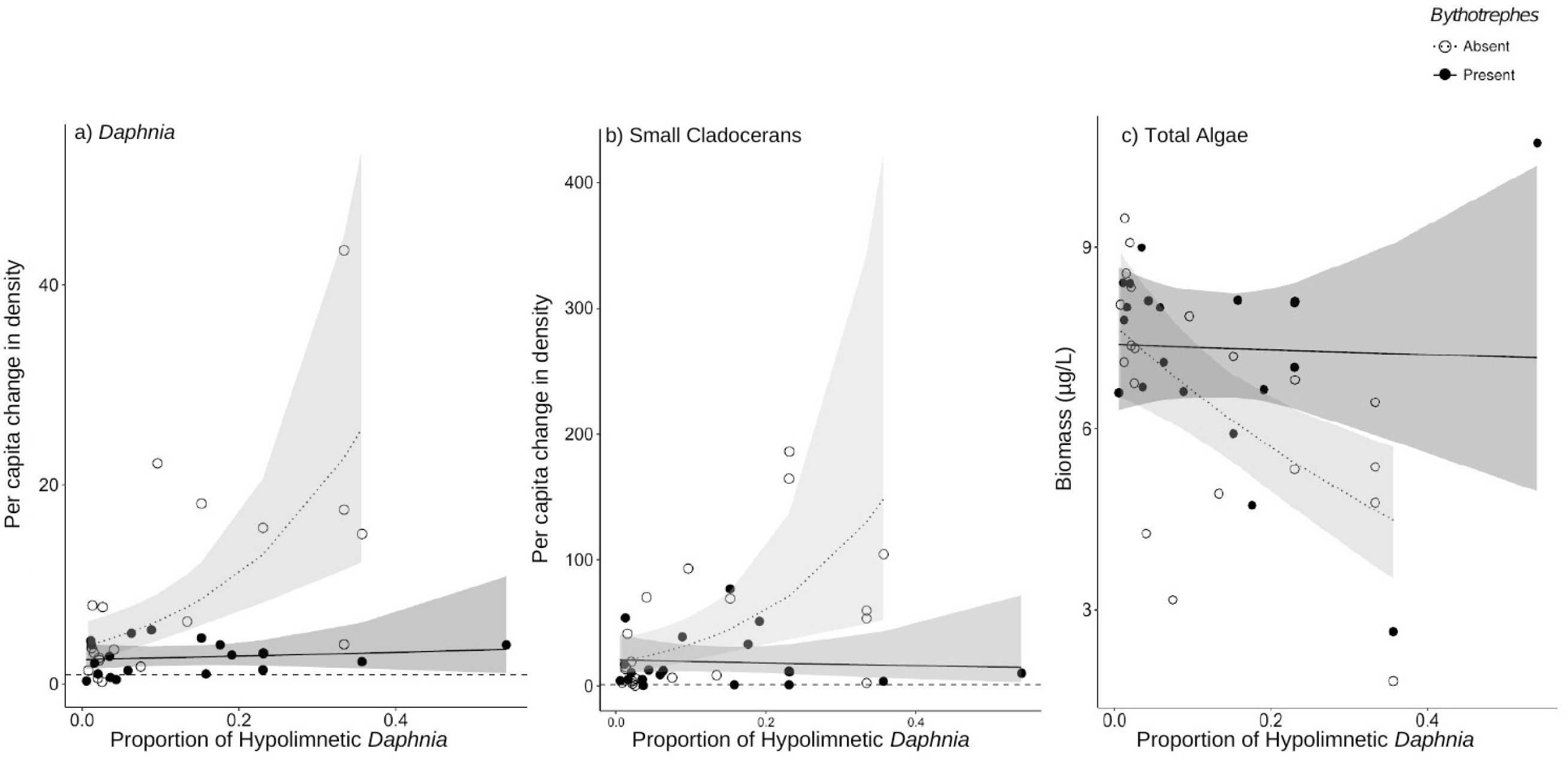
Effect off proportion of total hypolimnetic *Daphnia* in week 0 on the per capita change in a) Total *Daphnia* and b) Small cladoceran density, and c) Total algal biomass in week 3 in *Bythotrephes* absent and present mesocosms. Values above the dashed line at 1 indicate increasing density between week 0 and 3. Shaded regions represent the 95% confidence interval estimated from the best fitting model.

Small cladocerans were mostly epilimnetic and increased in density across all mesocosms during the experiment. We observed a significant interaction between *Bythotrephes* presence and the proportion of total hypolimnetic *Daphnia* in week 0 on per capita change in small cladoceran density (piecewise SEM, Figure 2a, Table S5). In uninvaded mesocosms, larger per capita increases in total small cladoceran density were observed in mesocosms with a greater proportion of total hypolimnetic *Daphnia* in week 0 (Gamma GLM, Figure 3b, p = 0.02, df = 36, Table S13). Total *Daphnia* depth distribution influenced the magnitude of *Bythotrephes* impact on total small cladocerans, with a smaller per capita increase in density observed in invaded mesocosms with a greater proportion of hypolimnetic *Daphnia,* as compared to uninvaded mesocosms (Figure 3b).

### Algal Biomass

We observed a significant interaction between *Bythotrephes* presence and proportion of hypolimnetic *Daphnia* in week 0 on total algal biomass at the end of study (Gamma Robust Regression, p = 0.0002, df = 37, Table S13). Total algal biomass was greater in invaded mesocosms as compared to uninvaded mesocosms, with the greatest increase observed in mesocosms with greatest proportion of hypolimnetic *Daphnia* in week 0 (Figure 3c, Table S13). Cyanobacteria were the most abundant functional group, followed by green algae, diatoms, and cryptophytes. Green algal biomass was predicted by the proportion of total *Daphnia* in the hypolimnion, and per capita change in density of large cladocerans, and juvenile copepods (Figure 2c, Vertical position SE: 0.44, Large cladocerans SE: 0.43, Juvenile copepods SE: 0.14, Table S6), and was greater in mesocosms with a greater proportion of hypolimnetic *Daphnia* in week 0 (Figure 4e). Diatom biomass was predicted by *Bythotrephes* presence, the proportion of hypolimnetic *Daphnia*, and per capita change in density of small cladocerans, large cladocerans, calanoids and juvenile copepods (Figure 2d, Table S7, *Bythotrephes* SE: -0.44, Vertical position SE:1.48, Small cladoceran SE: -0.62, Large cladoceran SE: 0.65, Calanoid SE: -1.75, Juvenile copepods SE: -0.28), and was greater in mesocosms with a greater proportion of hypolimnetic *Daphnia* (Figure 4f). There was no effect of *Bythotrephes* presence, the proportion of total hypolimnetic *Daphnia*, or per capita change in density in any zooplankton group on cyanobacteria and cryptophyta biomass (Tables S8-S9).

**Figure 4:**
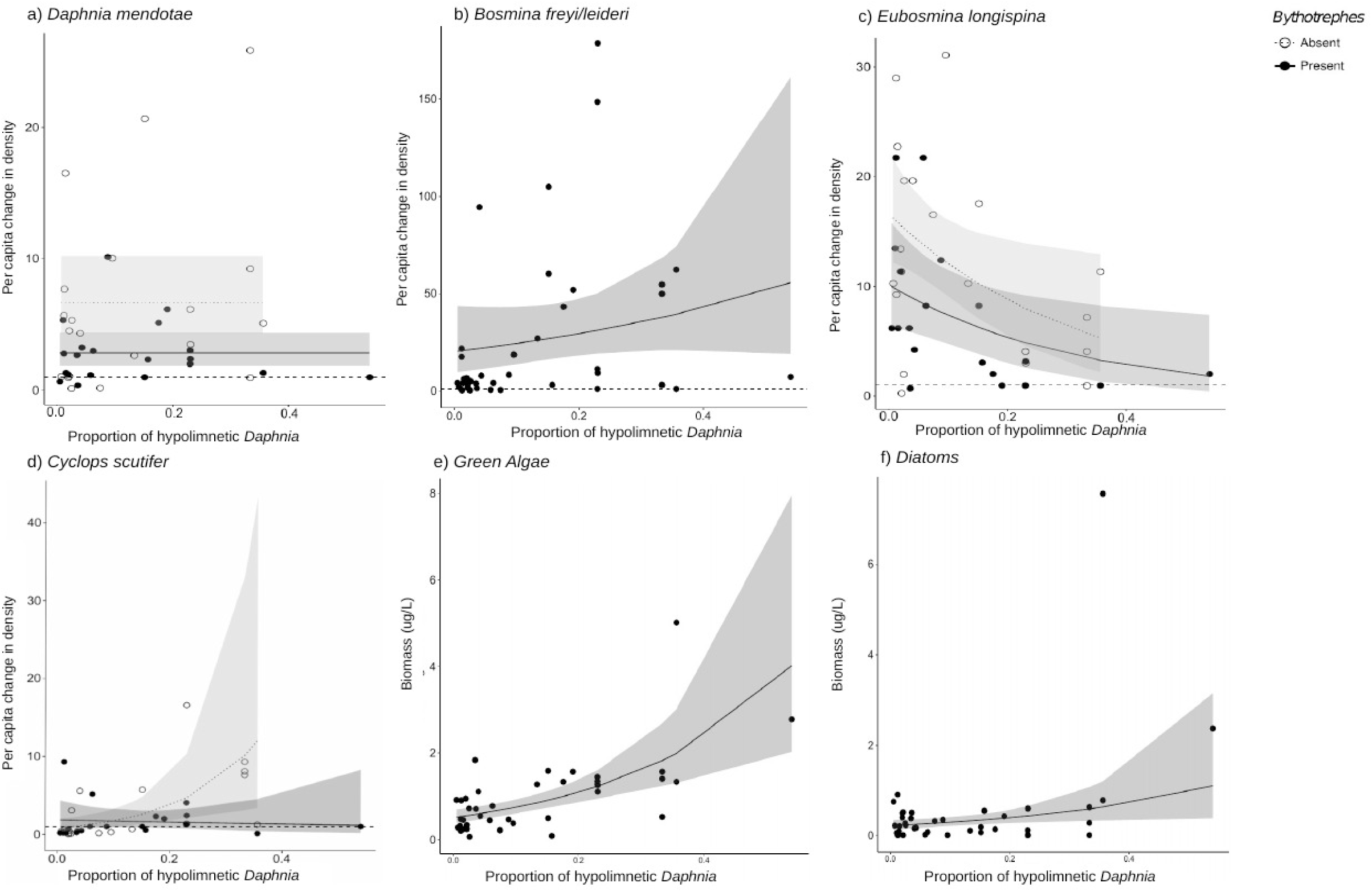
Effect of proportion of total hypolimnetic *Daphnia* in week 0 on the per capita change in a) *D. mendotae,* b) *B. freyi/leideri*, c) *E. longispina*, d) *C. scutifer* density, and biomass of e) green algae, and f) Diatoms in week 3 in *Bythotrephes* absent and present mesocosms. For green algae, no effect of *Bythotrephes* presence was detected. Values above the dashed line at 1 indicate increasing density between week 0 and 3. Shaded regions represent the 95% confidence interval estimated from the best fitting model.

### Species-level responses

For *Daphnia mendotae*, *Bythotrephes* presence predicted per capita change in density (piecewise SEM, Figure 2b, Table S10, *Bythotrephes* SE: -0.09). Per capita density for *D. mendotae* was lower in invaded mesocosms as compared to uninvaded mesocosms (Figure 4a). There was no effect of *Bythotrephes* presence or proportion of total hypolimnetic *Daphnia* on per capita change in *D. catawba* density (Table S10). Per capita change in density in *B. freyi/leideri* was only predicted by the proportion of total hypolimnetic *Daphnia* in week 0 (piecewise SEM, Figure 2b, Table S10, Vertical position SE: 0.01), with no effect of *Bythotrephes* presence. Increase in *B. freyi/*liederi density was greater in mesocosms with greater proportion of total hypolimnetic *Daphnia* (Figure 4b). Both *Bythotrephes* presence and proportion of total hypolimnetic *Daphnia* in week 0 predicted per capita change in *E. longispina* density (piecewise SEM, Figure 2b, Table S10, *Bythotrephes* SE: -0.03, Vertical position SE: -0.05). There was a smaller increase in *E. longispina* density (Figure 4c) in mesocosms with a greater proportion of total hypolimnetic *Daphnia* in week 0. There was no effect of *Bythotrephes* presence or the proportion of total hypolimnetic *Daphnia* in week 0 on per capita change in *E. tubicen* density (piecewise SEM, Table S10).

For copepods, per capita change in *C. scutifer* (cyclopoid) density was predicted by the proportion of total hypolimnetic *Daphnia* in week 0 and *Bythotrephes* presence (piecewise SEM, Figure 2b, Table S8, *Bythotrephes* SE: -0.18, Vertical position SE: 0.13). Per capita increase in *C. scutifer* density was smaller in invaded mesocosms as compared to uninvaded mesocosms with a greater proportion of hypolimnetic *Daphnia* (Figure 4d, Gamma GLMs, Table S13, p = 0.02, df = 36). There was no effect of *Bythotrephes* presence or proportion of total hypolimnetic *Daphnia* on per capita change in *S. oregonensis* (calanoid) density (Table S10).

## Discussion

Differences in the vertical position of *Daphnia* in our mesocosms resulted in differences in the effect of *Bythotrephes* predation on the change in abundance of small cladocerans as well as individual species (e.g., *C. scutifer*). Most notably, we found larger increases in phytoplankton biomass in invaded mesocosms where there was little spatial overlap between *Daphnia* and *Bythotrephes,* i.e., greater proportion of hypolimnetic *Daphnia* as compared to uninvaded mesocosms. The magnitude of increase was associated with the degree of spatial overlap. Larger increases in phytoplankton biomass were observed in invaded mesocosms with more hypolimnetic *Daphnia*. This confirms our hypothesis that reduced grazing due to *Bythotrephes* predation on small cladocerans and deeper *Daphnia* vertical position increase total algal biomass. Trophic cascades associated with *Bythotrephes* invasion are attributed to reduced *Daphnia* and cladoceran grazing due to *Bythotrephes* predation (Strecker and Arnott, 2008; Walsh et al., 2016). Our results suggest that differences in *Daphnia* vertical position can also contribute to trophic cascades in *Bythotrephes-*invaded systems. It is not known if differences in *Daphnia* vertical position can lead to different long-term community and ecosystem outcomes in *Bythotrephes* invaded lakes. While our results show clear differences in community level impacts, these were only observed in over a three-week period and we did not examine temporal dynamics over longer time scales.

Total *Daphnia* vertical position did not change during the experiment and did not influence *Bythotrephes* impact on total *Daphnia* abundance. However, *Bythotrephes* preferential predation on *Daphnia* in invaded mesocosms (Schulz and Yurista, 1999), resulted in smaller increases in abundances as compared to uninvaded mesocosms, matching observations from field surveys and mesocosm experiments (Strecker and Arnott, 2005; Pangle and Peacor, 2006; Strecker and Arnott 2008; Pangle and Peacor, 2009; Strecker and Arnott, 2010; Jokela et al., 2013; Azan and Arnott, 2017). This lack of *Daphnia* vertical position impact on total *Daphnia* abundance in invaded mesocosms could be due to similarity between the impacts of *Bythotrephes* predation and the metabolic costs of occupying deeper colder hypolimnetic waters (Kerfoot, 1985; Dawidowicz and Loose, 1992; Loose and Dawidowicz, 1994; Cole et al., 2002; Pangle et al., 2007; Pangle and Peacor, 2010). The vertical position of the most common *Daphnia* species, *D. mendotae* and *D. catawba,* did not change during the experiment (Table S10). Increase in *D. mendotae* abundance in invaded mesocosms was less as compared to uninvaded mesocosms, suggesting *Bythotrephes* predation on epilimnetic individuals. There was no effect of *Bythotrephes* on *D. catawba* density indicating a possible preferential *Bythotrephes* predation on larger bodied *Daphnia* (Schulz and Yuritsa, 1999), *D. mendotae* over the smaller *D. catawba*.

Using per capita change in density allowed us to directly compare between *Bythotrephes* treatments regardless of differences in starting densities among our mesocosms. However, we recognize that this metric is sensitive to sampling and demographic effects. We accounted for this by adding a minimum detection density to abundances in weeks 0 and 3 for all taxa assessed prior to any analysis. This reduced the sensitivity of per capita change in density values to large changes by reducing their magnitude, especially for taxa with low abundances at the start of the experiment and high abundances at the end of the experiment, such as small cladocerans(Tables S2-3). Low initial small cladoceran densities may be one driver for the large per capita increase we observed for this taxa regardless of *Bythotrephes* presence, although the observed coefficient of correlation between per capita change in small cladoceran density and initial density was not significantly different from bootstrapped values (Figure S6).

### Indirect impacts on other zooplankton groups

Fewer small cladocerans were present in invaded mesocosms with deeper *Daphnia* vertical position (i.e., greater proportion of hypolimnetic *Daphnia*). This may be linked to *Bythotrephes* preference for large bodied cladocerans such as *Daphnia* (Schulz and Yurista, 1999) in mesocosms where more *Daphnia* are epilimnetic. In mesocosms with a deeper *Daphnia*, *Bythotrephes* predation on small cladocerans led to the smaller increases observed. *Bythotrephes* predation reduces small cladoceran abundance (Vanderplog et al., 1993; Yan and Pawson, 1997; Yan et al., 2001; Barbiero and Tuchman, 2004; Strecker et al., 2006; Strecker and Arnott, 2008; Kerfoot et al., 2016), although this is not consistently observed (Lehman and Caeceres, 1993; Barbiero and Tuchman, 2004; Strecker and Arnott, 2005; Hessen et al., 2011). The effect of *Bythotrephes* predation on small cladocerans is varied, with declines in abundance ranging from 40-126% across invaded lakes (Vanderplog et al., 1993; Yan et al., 2001; Kerfoot et al., 2016). Our results suggest that *Daphnia* vertical position could be an important factor explaining this variation.

*B. freyi/leideri* density increased as the proportion of hypolimnetic *Daphnia* increased. For *E. longispina,* deeper *Daphnia* vertical position was associated with smaller increases in density. Furthermore, invaded mesocosms exhibited smaller *E. longispina* density increases as compared to uninvaded mesocosms. These contrasting effects suggest that *B. freyi/leideri* in our mesocosms was constrained by competition for grazing with *Daphnia* (Demott and Kerfoot, 1982). For *E. longispina*, *Bythotrephes* predation is likely underlying the smaller increase in density observed in invaded mesocosms. This contradicts observations from Kelly et al. (2012), where *E. longispina* abundance increased in *Bythotrephes*-invaded lakes, suggesting that *Bythotrephes* impacts may be mediated by other factors. It is unclear what is driving the lack of increase in *E. longispina* density with deeper *Daphnia* vertical position. A deeper *Daphnia* vertical position may increase competition between *E. longispina* and other bosminid species. However, competition between different bosminid species, especially in the presence and absence of *Daphnia* is unknown.

Contrary to our prediction, *Bythotrephes* presence did not impact change in total calanoid or *S. oregonensis* density. This is surprising as negative effects of *Bythotrephes* on calanoid copepods (Strecker and Arnott, 2005; Strecker et al., 2006; Hessen et al., 2011; Bourdeau et al., 2011; Kelly et al., 2012) have been broadly observed in the literature. Furthermore, the lack of an effect of deeper *Daphnia* vertical position on calanoid density also contradicts expected increases due to reduced competition between *Daphnia* and calanoids (Sommer et al., 2003). Both *Daphnia* and calanoids primarily feed on algae, therefore less competition for algae between *Daphnia* and calanoids is expected in mesocosms with greater proportion of hypolimnetic *Daphnia,* leading to an increase per capita calanoid density. We may not have detected the impacts of these competitive interactions due to the short experiment time frame.

We observed interactive negative effects of *Bythotrephes* and *Daphnia* vertical position on change in total cyclopoid density. A larger increase in *C. scutifer* density was observed in uninvaded mesocosms with more hypolimnetic *Daphnia* as compared to invaded mesocosms, suggesting that *Bythotrephes* presence negatively impacts *C. scutifer* density. Kelly et al. (2012) observed similar *Bythotrephes* impacts on *C. scutifer* abundance in Canadian and Norwegian lakes. In contrast, no *Bythotrephes* effect was observed by Hessen et al. (2011), while a positive effect was observed by Walseng et al. (2015). Although *Bythotrephes* effects on *C. scutifer* remain unresolved in observational studies, our results provide the first experimental evidence of negative *Bythotrephes* impacts on *C. scutifer* abundance. The negative effect of *Daphnia* vertical position on total cyclopoid and *C. scutifer* density may be a consequence of indirect interactions. Deeper *Daphnia* vertical position was linked to increased juvenile copepod density, which had a positive effect on total cyclopoid density. The negative effect of *Daphnia* vertical position on cyclopoids is likely not mediated through indirect effects on juvenile copepod density, and may be a result of competitive interactions which were not included in our analysis.

It is unclear why we observed changes in total *Daphnia* daytime vertical position in our mesocosms after inoculating them from lakes that have large differences in the proportion of hypolimnetic *Daphnia* (Echo Lake: 3% hypolimnetic, Bonnie Lake: 74% hypolimnetic). Daytime vertical position in *Daphnia* species is influenced by many ecological forces, including predator presence and location in the water column, predator type, food availability, exposure to ultra-violet radiation, and temperature-related metabolic costs (Leibold, 1990; Boeing et al., 2004; Williamson et al., 2011; Kessler & Lampert, 2004; Larsson & Lampert, 2012). In addition, there is also a strong genetic component, with large intra-population variation in vertical position observed in response to predator cues (De Meester 1993; De Meester 1996). Similar zooplanktivorous fish species were present in all three lakes (Ontario Ministry of Natural Resources and Forestry, 2024), making it unlikely that these changes were driven by different predator cues. Mesocosms were filled with water from Fletcher Lake, which has similar physicochemical properties as Echo and Bonnie lakes, with the exception of DOC, which was highest in Echo, followed by Fletcher, and Bonnie Lake (Fletcher: 4.6 mg/L, Bonnie: 2.3 mg/L, Echo: 6.6 mg/L). It is possible that differences in light availability and UV radiation as a consequence of differences in DOC influenced *Daphnia* vertical position in our mesocosms.

### Primary production and trophic cascades

*Daphnia* vertical position mediated the impacts of *Bythotrephes* on primary production by altering grazing pressure. Total algal biomass increased in invaded mesocosms with deeper *Daphnia* vertical position as compared to uninvaded mesocosms, likely a result of reduced small cladoceran abundance due to *Bythotrephes* predation and reduced epilimnetic grazing by *Daphnia*. The invasion of *Bythotrephes* in some North American temperate lakes is associated with trophic cascades due to reduced grazer biomass (Walsh et al., 2016; Martin et al., 2022) which has been observed in some lakes but not others (Strecker and Arnott, 2008). Our results suggest that in addition to the nutrient status of these lakes (Walsh et al., 2016), *Daphnia* vertical position could be an important factor explaining the varied *Bythotrephes* impacts on primary production in these studies.

Total algal biomass increased across most mesocosms (Figure S7) regardless of grazing by *Daphnia* and other cladocerans. *Daphnia* grazing was reduced in mesocosms with more hypolimnetic *Daphnia,* likely increasing green algal and diatom biomass. *Bythotrephes* presence negatively impacted diatom biomass. We also observed a negative effect of increasing small cladoceran density on diatom biomass. Apparent competition due to *Bythotrephes* preferential predation on larger-bodied *Daphnia* could lead to increased small cladoceran grazing. Furthermore, bosminids (i.e., *B. freyi/leideri*, *E. longispina*) were the most abundant small cladoceran taxa in our mesocosms. Their demonstrated selectivity on diatoms and green algae (Fulton III, 1988; Tonno et al., 2016) likely resulted in the decline observed.

## Conclusions

Food web and ecosystem consequences of differences in *Daphnia* vertical position remain largely ignored, despite literature suggesting that inter-population trait variation, especially in predator-prey interactions, may be the primary mechanism maintaining food web complexity in ecological systems and driving trophic cascades. This study highlights the strong influence of *Daphnia* vertical position on interactions between zooplankton groups, ultimately affecting primary production in lake ecosystems, regardless of *Bythotrephes* presence. Our results also provide the first experimental evidence suggesting that differences in *Daphnia* depth distribution influence the impacts of *Bythotrephes* predation on other cladoceran groups, resulting in increased algal biomass. Understanding the influence of *Daphnia* vertical position on the structure and functioning of lake ecosystems will improve our ability to predict impacts of future invasions.

## Acknowledgements

We thank the Dorset Environmental Science Centre (DESC) and its staff, especially James Rusak, Ron Ingram and Tim Field for providing logistical support during mesocosm sampling. We thank Sarah Lamb, Matthew Laird, and Shakira Azan for their field assistance.

## Data archiving statement

All data used in the analysis and presentation of results will be made available on Dryad at the time of publication.

## References

Azan, S. S. E., Arnott, S. E. and Yan N. D. (2015) A review of the effects of *Bythotrephes longimanus* and calcium decline on zooplankton communities — can interactive effects be predicted? Environ. Rev., 23, 395–413.

Azan, S. S. E., and Arnott, S.E. (2017) The effects of *Bythotrephes longimanus* and calcium decline on crustacean zooplankton communities in Canadian Shield lakes. Hydrobiologia, 785, 307–325.

Banks, P. and Dickman, C. R. (2007) Alien predation and the effects of multiple levels of prey naiveté. Trends Ecol. Evol., 22, 229–230.

Barbiero, R.P. and Tuchman, M. L. (2004) Changes in the crustacean communities of Lakes Michigan, Huron, and Erie following the invasion of the predatory cladoceran *Bythotrephes longimanus*. Can. J. Fish Aquat. Sci., 61, 2111–2125.

Barnett, A. J., Finlay, K. and Beisner, B. E. (2007) Functional diversity of crustacean zooplankton communities: towards a trait-based classification. Freshw. Biol., 52, 796–813.

Benard, M. F. (2004) Predator-induced phenotypic plasticity in organisms with complex life histories. Annu. Rev. Ecol. Syst., 35, 651–673.

Benesh, D. P. and Kalbe, M. (2016) Experimental parasite community ecology: intraspecific variation in a large tapeworm affects community assembly. J. Anim. Ecol., 85, 1004–1013.

Boeing, W. J., D. M. Leech & C. E. Williamson, 2004. Damaging UV radiation and invertebrate predation: conflicting selective pressures for zooplankton vertical distribution in the water column of low DOC lakes. Oecologia 138: 603–612.

Boeing, W. J., Ramcharan, C. W. and Riessen, H. P. (2006) Clonal variation in depth distribution of *Daphnia pulex* in response to predator kairomones. Arch. Hydrobiol., 166, 241–260.

Bolnick, D.I., Amarasekare, P., Araújo, M.S., Bürger, R., Levine, J.M., Novak, M., Rudolf, V.H., Schreiber, S.J., Urban, M.C. and Vasseur, D.A., (2011) Why intraspecific trait variation matters in community ecology. Trends Ecol. Evol., 26, 183–192.

Boudreau, S. A. and Yan, N. D. (2003) The differing crustacean zooplankton communities of Canadian Shield lakes with and without the nonindigenous zooplanktivore *Bythotrephes longimanus*. Canadian Journal of Fisheries and Aquatic Sciences, 60, 1307–1313.

Boudreau, S.A. and Yan, N. D. (2004) Auditing the accuracy of a volunteer based surveillance program for an aquatic invader, *Bythotrephes*. Environ. Monit. Assess., 91, 17–26.

Bourdeau, P. E., Pangle, K.L. and Peacor, S. D. (2011) The invasive predator *Bythotrephes* induces changes in the vertical distribution of native copepods in Lake Michigan. Biol. Invas., 13, 2533–2545.

Bourdeau, P. E., Pangle, K. L., Reed, E. M. and Peacor, S. D. (2013) Finely tuned response of native prey to an invasive predator in a freshwater system. Ecology, 94, 1449–1455.

Cairns, A., Yan, N.D., Weisz, E., Petruniak, J. and J. Hoare, J, (2007) Operationalizing CAISN project 1.V, Technical Report No. 2: the large inland lake *Bythotrephes* survey—limnology, database design, and presence of *Bythotrephes* in 311 south-central Ontario lakes. Dorset Environmental Science Centre, Dorset, Ontario, 66pp.

Cantoni, E., and Ronchetti, E. (2001). Robust inference for generalized linear models. J. Am. Stat. Assoc., 96, 1022–1030.

Cantoni, E., and E. Ronchetti, E. (2006). A robust approach for skewed and heavy-tailed outcomes in the analysis of health care expenditures. J Health Econ., 25,198–213.

Carpenter, S. R., Kitchell, J. F. and Hodgson, J. R. (1985) Cascading trophic interactions and lake productivity. BioScience, 35, 634–639.

Carthey, A. J. R. and Banks, P. B. (2014) Naïveté in novel ecological interactions: lessons from theory and experimental evidence. Biol. Rev., 89, 932–949.

Carthey, A. J. R., Bucknall, M. P., Wierucka, K. and Banks, P. B. (2017) Novel predators emit novel cues: a mechanism for prey naivety towards alien predators. Sci. Rep., 7, 16377.

Cole, P. C., Luecke, C., Wurtsbaugh, W. A. and Burkart, G. (2002) Growth and survival of *Daphnia* in epilimnetic and metalimnetic water from oligotrophic lakes: the effects of food and temperature. Freshw. Biol., 47, 2113–2122.

Cousyn, C., De Meester, L., Colbourne, J.K., Brendock, L., Verschuren, D. and Volkaert, F. (2001). Rapid, local adaptation of zooplankton behavior to changes in predation pressure in the absence of neutral genetic changes. Proc. Natl. Acad. Sci., 98, 6256–6260.

Cox, J. G. and Lima, S. L. (2006) Naiveté and an aquatic–terrestrial dichotomy in the effects of introduced predators. Trends Ecol. Evol., 21, 674–680.

Crispo, E., (2008). Modifying effects of phenotypic plasticity on interactions among natural selection, adaptation and gene flow. J. Evol. Biol., 21, 1460–1469.

Crawley, M. J. (2011) Statistics: An introduction using R. John Wiley and Son Ltd.

Dawidowicz, P. and Loose, C.J. (1992) Metabolic costs during predator-induced diel vertical migration of *Daphnia*. Limnol. Oceanogr., 37, 1589–1595.

De Meester, L. (1993) Genotype, fish mediated chemical, and phototactic behavior in *Daphnia magna*. Ecology, 74, 1467–1474.

De Meester, L. (1996) Evolutionary potential and local genetic differentiation in a phenotypically plastic trait of a cyclical parthenogen, *Daphnia magna*. Evolution 50: 1293–1298.

Melo, R.D. and Hebert, P. D. N. (1994) A taxonomic reevaluation of North American Bosminidae. Can. J. Zool., 72, 1808–1825.

DeMott, W. R. and Kerfoot, W. C. (1982) Competition Among cladocerans: Nature of the interaction between *Bosmina* and *Daphnia*. Ecology, 63, 1949–1966.

Dieckmann, U., Doebeli, M., Metz, J. A. J., and Tautz. D., (2004). Adaptive speciation. Cambridge University Press

Elton, C., (1958) The ecology of invasions by animals and plants. University of Chicago Press

Finlay, K. and Roff, J. C. (2004) Radiotracer determination of the diet of calanoid copepod nauplii and copepodites in a temperate estuary. J. Mar. Syst., 61, 552–562.

Fulton, R.S., III (1988) Grazing on filamentous algae by herbivorous zooplankton. Freshw. Biol., 20, 263–271.

Haney, J. F., M. A. Aliberti, E. Allan, S. Allard, D. J. Bauer, W. Beagen, S. R. Bradt, B. Carlson, S. C. Carlson, U. M. Doan, J. Dufresne, W. T. Godkin, S. Greene, A. Kaplan, E. Maroni, S. Melillo, A. L. Murby, J. L. Smith, B. Ortman, J. E. Quist, S. Reed, T. Rowin, M. Schmuck, R. S. Stemberger and Travers, B. (2013) An-Image-based Key to the Zooplankton of North America, version 5.0 released 2013. University of New Hampshire Center for Freshwater Biology. http://cfb.unh.edu.

Hessen, D. O., Bakkestuen, V. and Walseng, B. (2011) The ecological niches of *Bythotrephes* and *Leptodora*: Lessons for predicting long-term effects of invasion. Biol. Invas., 13, 2561–2572.

Ibarra-Isassi, J., Handa, I. T., and Lessard, J.P. (2022) Community-wide trait adaptation, but not plasticity, explains ant community structure in extreme environments. Funct. Ecol., 37, 139–149.

Jokela, A., Arnott, S.E., and Beisner, B. E. (2011) Patterns of *Bythotrephes longimanus* distribution relative to native macroinvertebrates and zooplankton prey. Biol. Invas., 13, 2573–2594.

Jokela, A. M., Arnott, S. E. and Beisner, B. E. (2013) Influence of light on the foraging impact of an introduced predatory cladoceran, *Bythotrephes longimanus*. Freshw. Biol., 58, 1946–1957.

Jokela, A. M., S. E. Arnott, and Beisner, B. E. (2017) Biotic resistance of impact: a native predator (*Chaoborus*) influences the impact of an invasive predator (*Bythotrephes*) in temperate lakes. Biol. Invas., 19, 1495–1515.

Kelly, N.E., Yan, N.D., Walseng, B. and Hessen, D. O. (2012) Differential short- and long-term effects of an invertebrate predator on zooplankton communities in invaded and native lakes. Divers. Distrib., 19, 396–410.

Kerfoot, W. C., (1985) Adaptive value of vertical migration: comments on the predation hypothesis and some alternatives. Contributions in Marine Science Supplementary, 27, 91–113.

Kerfoot, W. C., Hobmeier, M. M., Yousef, F., Lafrancois, B. M., Maki, R. P. and Hirsch, J. K. (2016). A plague of waterfleas (*Bythotrephes*): impacts on microcrustacean community structure, seasonal biomass, and secondary production in a large inland-lake complex. Biol. Invas., 18, 1121–1145.

Korovchinsky, N. M. and Arnott, S. E. (2019). Taxonomic resolution of the North American invasive species of the genus *Bythotrephes* Leydig, 1860 (Crustacea: Cladocera: Cercopagididae). Zootaxa, 4691, 125–138.

Larsson, P. and Lampert, W. (2012). Finding the optimal vertical distribution: Behavioural responses of *Daphnia pulicaria* to gradients of environmental factors and the presence of fish. Freshw. Biol., 57, 2514–2525.

Lehman, J. T. and Cáceres, C. E. (1993). Food-web responses to species invasion by a predatory invertebrate: *Bythotrephes* in Lake Michigan. Limnol. Oceanogr., 38, 879–891.

Lehman, J. T., Bilkovic, D. M. and Sullivan, C. (1997) Predicting development, metabolism and secondary production for the invertebrate predator *Bythotrephes*. Freshw. Biol., 38, 343–352.

Leibold, M. A. (1990). Resources and predators can affect the vertical distributions of zooplankton. Limnol. Oceanogr., 35, 938–944.

Lenhart, P. A., Jackson, K. A. and White, J. A. (2018) Heritable variation in prey defence provides refuge for subdominant predators. Proc. R. Soc. B., 285, 20180523.

Lima, S. I. (1998). Nonlethal effects in the ecology of predator-prey interactions: what are the ecological effects of anti-predator decision-making? BioScience, 48, 25–34.

Loose, C. J. and Dawidowicz, P. (1994). Trade-offs in diel vertical migration by zooplankton: The costs of predator avoidance. Ecology, 75, 2255–2263.

Mack, R. N., Simberloff, D., Lonsdale, W. M., Evans, H., Clout, M. and Bazzaz, F. A. (2000) Biotic invasions: causes, epidimiology, global consequences, and control. Ecol. Appl., 10,689–710.

Martin, B.E., Walsh, J.R. &Vander Zanden, J.M. (2022) Rise of a native apex predator and an invasive zooplankton cause successive ecological regime shifts in a North Temperate Lake. Limnol. Oceanogr., 67, S163–S172.

Morris, D. P., H. Zagarese & C. E. Williamson (1995) The attenuation of solar UV radiation in lakes and the role of dissolved organic carbon. Limnology and Oceanography 40: 1381–1391.

NASA Langley Atmospheric Science Data Center, 2016. Surface Meterology and Solar Energy: a renewable energy resource website (release 6.0). Accessed June 1, 2016, from http://power.larc.nasa.gov/cgi-bin/cgiwrap/solar/hirestimeser.cgi?

Ontario Ministry of Natural Resources and Forestry (2020) Lake Fact Sheets. Accessed April 1^st^ 2024 from http://www.muskokawaterweb.ca/lake-data/mnr-data/lake-fact-sheet

Pangle, K. L. and Peacor, S.D. (2006) Non-lethal effect of the invasive predator *Bythotrephes longimanus* on *Daphnia mendotae*. Freshw. Biol., 51, 1070–1078.

Pangle, K. L. and Peacor, S. D. (2009) Light dependent by the invertebrate planktivore *Bythotrephes longimanus*. Can. J. Fish Aquat. Sci., 66, 1748–1757.

Pangle, K. L., Peacor, S. D., Johannsson, O. E. and Field, E. (2007) Large nonlethal effects of an invasive invertebrate predator on zooplankton population growth rate. Ecology, 88, 402–412.

Paine, R. T. (1966) Food web complexity and species diversity. Amer. Nat., 910, 65–75.

Paine, R. T. (1980) Food webs: linkage, interaction strength, and community infrastructure. J. Anim. Ecol., 49, 667–685.

Palkovacs, E. P. and Post, D. M. (2009) Experimental evidence that phenotypic divergence in predators drives community divergence in prey. Ecology, 90, 300–305.

Pfennig, D. W., Wund, M. A., Snell-Rood, E. C., Cruickshank, T., Schlichting, C. D., and Moczek, A. P. (2010) Phenotypic plasticity’s impacts on diversification and speciation. Trends Ecol. Evol., 25, 459–467.

Polis, G. A. and Holt, R. D. (1992) Intraguild predation: the dynamics of complex trophic interactions. Trends Ecol. Evol., 7, 151–154.

Post, D. M., Palkovacs, E. P., Schielke, E. G. and Dodson, S. I. (2008) Intraspecific variation in predator affects community structure and cascading trophic interaction. Ecology, 89, 2019–2032

Salo, T., Mattila, J. and Eklöf, J. (2020) Long-term warming affects ecosystem functioning through species turnover and intraspecific trait variation. Oikos, 129, 283–295

Sih, A., Bolnick, D.I., Luttbeg, B., Orrock, J.L., Peacor, S.D., Pintor, L.M., Preisser, E., Rehage, J.S., and Vonesh, J. R. (2010) Predator–prey naïveté, antipredator behavior, and the ecology of predator invasions. Oikos, 119, 610–621

Schmitz, O., Křivan, V. and Ovadia, O. (2004) Trophic cascades: the primacy of trait-mediated indirect interactions. Ecol Lett., 7, 153–163.

Schulz, K. L. and Yurista, P. M. (1999) Implications of an invertebrate predator’s (*Bythotrephes cederstroemi*) atypical effects on a pelagic zooplankton community. Hydrobiologia, 380, 179– 193.

Smith, K. and Fernando, C.H. (1978) A guide to the freshwater calanoid and cyclopoid copepod Crustacea of Ontario. University of Waterloo, Department of Biology. Ser. No. 18

Strecker, A.L. and Arnott, S. E. (2005) Impact of *Bythotrephes* invasion on zooplankton communities in acid-damaged and recovered on the Boreal Shield. Can. J. Fish Aquat. Sci., 62, 2450–2462.

Strecker, A.L., Arnott, S.E., Yan, N.D. and Girard, R. (2006) Variation in the response of crustacean zooplankton species richness and composition to the invasive predator *Bythotrephes longimanus*. Can. J. Fish Aquat. Sci., 63, 2126–2136.

Strecker, A.L. and Arnott, S. E. (2008) Invasive predator, *Bythotrephes*, has varied effects on ecosystem function in freshwater lakes. Ecosystems, 11, 490–503.

Sommer, U., Sommer, F., Santer, B., Zöllner, E., Jürgens, K., Jamieson, C., Boersma, M., and Gocke, K. (2003) *Daphnia* versus copepod impact on summer phytoplankton: functional compensation at both trophic levels. Oecologia, 135, 639–647.

Tessier, A. J. and Leibold, M.A. (1997) Habitat use and ecological specialization within lake *Daphnia* populations. Oecologia, 109, 561–570.

Tollrian, R. and Harvell, C. D. (1999) The ecology and evolution of inducible defenses. Princeton University Press

Tõnno, I., Agasild, H., Kõiv, T., Freiberg, R., Nõges, P., and Nõges, T. (2016). Algal diet of small-bodied crustacean zooplankton in a cyanobacteria-dominated eutrophic lake. PloS one, 11, e0154526

Trussell, G. C., Ewanchuk, P. J. and Bertness, M. D. (2003) Trait-mediated effects in rocky intertidal food chains: predator risk cues alter prey feeding rates. Ecology, 84, 629–640.

Vanderploeg, H.A., Liebig, J.R. and Omair, M. (1993) *Bythotrephes* predation on Great Lakes zooplankton measured by an in-situ method implication for zooplankton community structure. Arch. Hydrobiol., 127, 1–8.

Vanni, M. J. (1986) Competition in zooplankton communities: Suppression of small species by *Daphnia pulex*. Limnol. Oceanogr., 31, 1039–1056.

Walsh, J. R., Carpenter, S. R. and Vander Zanden, M. (2016) Invasive species triggers a massive loss of ecosystem services through a trophic cascade. Proc. Natl. Acad. Sci., 113, 4081–4085.

Walseng, B., Andersen, T., and Hessen, D. O. (2015) Higher zooplankton species richness associated with an invertebrate top predator. Freshw. Biol., 60, 903–910.

Ward, H. B. and Whipple, G. C. (1959) In Edmondson, W. T. (eds), Fresh Water Biology, 2nd edn. Wiley,

Watt, P. J. and Young, S. (1994) Effect of predator chemical cues on *Daphnia* behavior in both horizontal and vertical planes. Animal Behaviour, 48, 861–869.

Williamson, C. E., 2009. UV Lakes project: Water Transparency and UV attenuation. Accessed June 1, 2016, from http://www.orgs.miamioh.edu/uvlakes/UVecology/Intro/intro.html.

Williamson, C. E., Fischer, J. M., Bollens, S. M., Overholt, E. P. and Breckenridge, J. K. (2011) Toward a more comprehensive theory of zooplankton diel vertical migration: Integrating ultraviolet radiation and water transparency into the biotic paradigm. Limnol. Oceanogr., 56, 1603–1623.

Wissel, B. and Ramacharan, C. W. (2003) Plasticity of vertical distribution of crustacean zooplankton in lakes with varying levels of water colour. J. Plankt. Res., 25, 1047–1057.

Witty, L. M. (2004). Practical Guide to Identifying Freshwater Crustacean Zooplankton. Laurentian University, Cooperative Freshwater Ecology Unit, Department of Biology, Sudbury: 57 pp.

Yan, N. D. and Pawson, T. W. (1997) Seasonal variation in the size and abundance of the invading *Bythotrephes* in Harp Lake, Ontario, Canada. Hydrobiologia, 8, 157–168.

Yan, N. D., Blukacz, A., Sprules, W. G., Kindy, P. K., Hackett, D., Girard, R. D. and Clark, B. J. (2001) Changes in zooplankton and the phenology of the spiny water flea, *Bythotrephes*, following its invasion of Harp Lake, Ontario, Canada. Can. J. Fish Aquat. Sci., 58, 2341–2350.

