## Supplementary Materials for "Variation in behaviour of native prey mediates the impact of an invasive species on plankton communities"

#### Minimum detection densities

To account for the possibility of non-detections, we added a number representing the minimum density that could be detected (1 individual /total volume of mesocosm sampled in a single vertical tow) to densities of all taxa assessed in samples from week 0 and week 3. We added one third of the minimum detection density to each layer to represent the presence of a single undetected individual in that layer, for each species. We also added a minimum detection biomass (0.01 µg/L; representing the measurement resolution of the algae analyzer that we used) to all algal groups as it was unclear if a biomass of 0 represented a true absence, a lack of detection due to sampling, or measurement error (Table S4 provides number of mesocosms where minimum detection density was added for each major taxonomic group assessed). There is support for this protocol because we detected individuals of major taxonomic groups at the end of experiment in many mesocosms without detections in week 0 (Table S4).

#### *Daphnia* vertical position

After zooplankton stocking, the vertical position of *Daphnia* prior to the start of the experiment did not reflect the position in their source lakes. Instead, stocking mesocosms from these uninvaded lakes with different *Daphnia* vertical positions created a gradient of daphniid vertical position from 0 to 40% hypolimnetic (Figure S1).

Through the course of the experiment, it was possible that *Daphnia* could change its vertical position in response to individual mesocosm environments. For invaded mesocosms with greater spatial overlap between *Daphnia* and *Bythotrephes* (fewer hypolimnetic *Daphnia*), *Daphnia* may adopt a deeper depth distribution in response to *Bythotrephes* kairomone (Bourdeau et al. 2013). Additionally, in uninvaded mesocosms with more hypolimnetic *Daphnia*, metabolic costs associated with residing in deeper, colder waters with potentially lower algal biomass and quality could result in *Daphnia* migrating to warmer, algae abundant epilimnetic waters (Johnsen and Jakobsen 1987, Leibold 1990, Rautio et al. 2003, Kessler and Lampert 2004).

To determine if total *Daphnia* and individual *Daphnia* species shifted their depth distributions during the experiments, either as a result of direct predation or an induced response to *Bythotrephes* predation, I assessed differences in the proportion of hypolimnetic individuals with experiment week (categorical variables with two levels; Week 0 or Week 3) and *Bythotrephes* (categorical variables with two levels; Present or Absent) as interacting explanatory variables using either gamma or log-normal distributed generalized linear mixed models (GLMMs) with a log link and mesocosm as a random effect. Minimum adequate models were chosen using log-likelihood ratio tests based on Crawley's (2005) procedure. Any changes in the proportion of hypolimnetic *Daphnia* between week 0 and week 3 were interpreted as shifts in vertical position during the experiment. AICc values were used to assess model fit between log-normal and gamma distributed models, with the model with the lowest value chosen. The inclusion of mesocosm as a random effect was determined on the basis of log likelihood ratio tests using Crawley's (2008) procedure. Generalized linear hypothesis tests performed (GLHT) for relevant comparisons when significant interactions between predictor variables were detected. A summary of the results from this analysis is provided in Table S12. Figures S2 and S3 show vertical position for total *Daphnia, D. catawba, and D. mendotae* at the start of the experiment. Differences in vertical position for *D. catawba*, and *D. mendotae* between Week 0 and Week 3 are shown in figure S3.

**Zooplankton species categorization**

*Bythotrephes* negatively impacts cladoceran density (Boudreau & Yan, 2003; Strecker et al., 2006) and preferentially feeds on larger bodied cladocerans, especially *Daphnia* (Schulz & Yurista, 1999). No cladocerans between 0.85-1.0mm were found in our mesocosms. Non-*Daphnia* cladoceran species were categorized as small or large based on average species body length (Barnett et al. 2007). Small cladocerans included *Bosmina freyi/liederi, Eubosmina tubicen, Eubosmina longispina, Chydorus sphaericus, Eubosmina coregoni, Ceriodaphnia lacustris, Simocephalus vetulus,* and *Scapholeberis mucronata*. Large cladocerans included *Holopedium glacialis* and *Sida crystallina*. All of these species are impacted by *Bythotrephes* presence (Azan et al., 2015). Copepod zooplankton were split into calanoid copepods, cyclopoid copepods, nauplii and copepodites, representing different trophic positions and feeding strategies (calanoid and cyclopoid copepods), and juvenile stages (copepodites and nauplii). Calanoid copepods are suspension feeding herbivore-omnivores that could compete with *Daphnia* and other cladocerans for algal resources while cyclopoid copepods are raptorial feeding omnivore-carnivores (Barnett et al., 2007), that could be indirectly impacted by *Bythotrephes* predation on small cladocerans.

**Structural Equation Models**

For the full model, we only fit paths between *Bythotrephes* presence, proportion of total hypolimnetic *Daphnia* in week 0, and the per capita change in density for each zooplankton group (Figure 1). Paths for calanoid and cyclopoid copepods also included per capita change in copepodite density as an explanatory variable since copepodites mature into adult calanoid and cyclopoid copepods, directly impacting the per capita change in density for these taxonomic groups. Paths for total algal density included per capita change in density for all herbivorous zooplankton groups, i.e., *Daphnia,* all cladocerans, copepodites, and calanoid copepods as the density of these groups directly reduce total algal density across time. Collinearity between per capita change in density of all taxonomic groups was accounted for by explicitly treating these as correlated errors (not a directed path). Proportion of total hypolimnetic *Daphnia* in week 0 and *Bythotrephes* presence were included as exogenous variables and potential paths were fit with GLMs (normal distribution, log link function). Paths for per capita changes in calanoid and cyclopoid densities included per capita change in copepodite density as an exogenous variable. For total algal density, per capita change in densities for all herbivorous zooplankton groups were also fit as exogenous variables. We fit a nested version containing only the significant links identified in the full model to confirm, using AIC, that the full model is not more likely than the reduced model, given the data.

**Density dependent effects of *Bythotrephes* predation due to differences in starting densities**

Due to differences in starting densities (Table S2), it is possible that any significant influence of *Daphnia* vertical position on *Bythotrephes* impacts for any zooplankton group could be confounded by density dependent *Bythotrephes* predation. To ensure that density dependent predation was not confounding *Bythotrephes* impacts, we examined the correlation between total zooplankton density in week 0 and per capita change in density for taxa where *Daphnia* vertical position was found to influence per capita density. As per capita density is calculated by dividing densities in week 3 by densities in week 0, we expected per capita density for each taxa assessed to be spuriously correlated with total zooplankton density in week 0. Therefore, we assessed if the Pearson correlation coefficient between taxon per capita change in density and total zooplankton density in week 0 obtained from this analysis was observed within 95% of the distribution of coefficients obtained by bootstrapping (with replacement) (n = 1000). For all zooplankton taxa, correlation coefficients were observed within the 95$^{th}$ percentile of the bootstrapped distribution suggesting that per capita change in density observed in these groups was not influenced by differences in starting densities across my mesocosms (Figure S4).

**Influence of vertical position of *D. mendotae* and *D. catawba* on per capita change in density for major zooplankton taxonomic groups**

Proportion of hypolimnetic *D. mendotae* and *D. catawba* in week 0 impacted per capita change in total *Daphnia* density, with an increase in per capita total *Daphnia* density observed in mesocosms with a greater proportion of *D. mendotae* and *D. catawba* in the hypolimnion (Table S6 & S7). Proportion of hypolimetic *D. catawba* in week 0 negatively impacted per capita cyclopoid density and positively impacted per capita total algal biomass (Table S7). There was no effect of proportion of hypolimnetic individuals of these two species in week 0 on per capita change for any other zooplankton taxonomic group.

### References


Azan, S. S. E., Arnott, S. E. and Yan N. D. (2015) A review of the effects of *Bythotrephes longimanus* and calcium decline on zooplankton communities can interactive effects be predicted? *Environ. Rev.*, **23**, 395–413

Barnett, A. J., Finlay, K. and Beisner, B. E. (2007) Functional diversity of crustacean zooplankton communities: towards a trait-based classification. *Freshw. Biol.,* **52**, 796-813

Bourdeau, P. E., Pangle, K. L., Reed, E. M. and Peacor, S. D. (2013) Finely tuned response of native prey to an invasive predator in a freshwater system. *Ecology*, **94**, 1449-1455.

Boudreau, S. A. and Yan, N. D. (2003) The differing crustacean zooplankton communities of Canadian Shield lakes with and without the nonindigenous zooplanktivore *Bythotrephes longimanus*. *Canadian Journal of Fisheries and Aquatic Sciences*, **60**, 1307-1313.

Crawley, M. J. (2011) Statistics: An introduction using R. John Wiley and Son Ltd

Johnsen, G. H., & Jakobsen, P. J. (1987). The effect of food limitation on vertical migration in Daphnia longispina 1. *Limnology and Oceanography*, **32**, 873-880.

Kessler, K., & Lampert, W. (2004). Depth distribution of Daphnia in response to a deep-water algal maximum: the effect of body size and temperature gradient. *Freshwater biology*, *49*(4), 392-401

Leibold, M. A. (1990). Resources and predators can affect the vertical distributions of zooplankton. Limnol. Oceanogr., **35**, 938–944

Rautio, M., Korhola, A., & Zellmer, I. D. (2003). Vertical distribution of *Daphnia longispina* in a shallow subarctic pond: Does the interaction of ultraviolet radiation and *Chaoborus* predation explain the pattern?. *Polar Biology*, **26**, 659-665.

Schulz, K. L. and Yurista, P. M. (1999) Implications of an invertebrate predator's (*Bythotrephes cederstroemi*) atypical effects on a pelagic zooplankton community. Hydrobiologia, **380**, 179–193.

Strecker, A.L., Arnott, S.E., Yan, N.D. and Girard, R. (2006) Variation in the response of crustacean zooplankton species richness and composition to the invasive predator *Bythotrephes longimanus. Can. J. Fish Aquat. Sci.*, **63**, 2126–2136.

### Tables

Table S1: Weekly temperature measurements (°C) for Fletcher lake for 1-14m at 1m intervals through the duration of the experiment

| Depth (m) | Week 0 | Week 1 | Week 2 | Week 3 |
|---|---|---|---|---|
| 1 | 21.89 | 22.6 | 20.6 | 21.0 |
| 2 | 21.82 | 22.6 | 20.5 | 21.0 |
| 3 | 21.41 | 22.6 | 20.5 | 21.0 |
| 4 | 19.54 | 19.3 | 20.4 | 21.0 |
| 5 | 14.89 | 13.6 | 14.8 | 20.2 |
| 6 | 9.49 | 9.3 | 10.1 | 15.5 |
| 7 | 7.32 | 7.7 | 7.9 | 11.4 |
| 8 | 6.31 | 6.8 | 6.5 | 9.6 |
| 9 | 5.74 | 5.9 | 5.7 | 8.3 |
| 10 | 5.39 | 5.6 | 5.3 | 7.1 |
| 11 | 5.08 | 5.3 | 5.1 | 6.5 |
| 12 | 4.98 | 5.0 | 4.9 | 6.1 |
| 13 | 4.89 | 4.9 | 4.8 | 5.8 |
| 14 | 4.79 | 4.8 | 4.7 | 5.4 |

Table S2: Initial densities (individuals per litre) in each mesocosm for *Daphnia*, small cladocerans, large cladocerans, calanoids, cyclopoids, copepodids, and nauplii prior to the application of the *Bythotrephes* treatment (N = absent, Y = present). Mean ± standard error (SE) for each taxonomic group is also provided. Densities for zooplankton taxa are provided in individuals per litre while total algal density is provided in micrograms per litre.

| Mesocosm Number | *Bythotrephes* Treatment | Stocking Lake | *Daphnia* | Small Cladocerans | Large Cladocerans | Copepodites | Nauplii | Calanoids | Cyclopoids | Total Algae |
|---|---|---|---|---|---|---|---|---|---|---|
| 1 | N | Mix | 0.009 | 0.009 | 0.009 | 0.345 | 0.028 | 0.009 | 0.028 | 6.530 |
| 2 | N | Echo | 0.869 | 0.074 | 0.112 | 2.346 | 0.308 | 0.084 | 0.317 | 6.190 |
| 3 | N | Bonnie | 0.009 | 0.018 | 0.046 | 0.906 | 0.074 | 0.028 | 0.074 | 6.410 |
| 4 | Y | Bonnie | 0.018 | 0.009 | 0.018 | 0.355 | 0.018 | 0.028 | 0.018 | 6.300 |
| 5 | Y | Bonnie | 0.028 | 0.028 | 0.177 | 0.803 | 0.037 | 0.037 | 0.037 | 6.590 |
| 6 | Y | Mix | 0.458 | 0.074 | 0.074 | 1.140 | 0.121 | 0.018 | 0.140 | 5.940 |
| 7 | N | Bonnie | 0.018 | 0.009 | 0.018 | 0.448 | 0.037 | 0.056 | 0.046 | 5.780 |
| 8 | Y | Echo | 0.187 | 0.018 | 0.028 | 0.551 | 0.037 | 0.046 | 0.037 | 6.020 |
| 9 | Y | Bonnie | 0.028 | 0.009 | 0.159 | 0.701 | 0.018 | 0.037 | 0.018 | 6.180 |
| 10 | N | Bonnie | 0.112 | 0.018 | 0.093 | 0.738 | 0.009 | 0.009 | 0.009 | 5.820 |
| 11 | N | Mix | 0.551 | 0.028 | 0.215 | 1.065 | 0.074 | 0.009 | 0.074 | 4.230 |
| 12 | N | Echo | 0.495 | 0.046 | 0.065 | 0.981 | 0.074 | 0.074 | 0.074 | 6.980 |
| 13 | Y | Echo | 0.205 | 0.009 | 0.018 | 1.102 | 0.168 | 0.046 | 0.168 | 6.410 |
| 14 | Y | Echo | 0.850 | 0.037 | 0.187 | 0.944 | 0.187 | 0.018 | 0.187 | 6.640 |
| 15 | N | Bonnie | 0.009 | 0.028 | 0.252 | 0.616 | 0.056 | 0.037 | 0.056 | 6.800 |
| 16 | Y | Bonnie | 0.018 | 0.018 | 0.130 | 0.551 | 0.065 | 0.018 | 0.065 | 6.600 |
| 17 | N | Bonnie | 0.028 | 0.037 | 0.224 | 1.168 | 0.140 | 0.065 | 0.140 | 6.880 |
| 18 | Y | Mix | 0.364 | 0.028 | 0.046 | 2.009 | 0.196 | 0.037 | 0.196 | 6.910 |
| 19 | N | Mix | 0.159 | 0.056 | 0.102 | 1.897 | 0.243 | 0.037 | 0.243 | 6.390 |
| 20 | N | Echo | 0.607 | 0.056 | 0.130 | 3.093 | 0.411 | 0.112 | 0.411 | 7.320 |
| 21 | Y | Bonnie | 0.018 | 0.018 | 0.074 | 0.654 | 0.093 | 0.037 | 0.093 | 7.010 |
| 22 | N | Echo | 0.803 | 0.018 | 0.028 | 2.383 | 0.205 | 0.187 | 0.205 | 6.240 |
| 23 | Y | Bonnie | 0.028 | 0.009 | 0.065 | 0.841 | 0.074 | 0.046 | 0.074 | 6.100 |
| 24 | Y | Echo | 0.355 | 0.009 | 0.018 | 1.598 | 0.168 | 0.093 | 0.168 | 6.820 |
| 25 | N | Echo | 0.102 | 0.009 | 0.084 | 0.831 | 0.448 | 0.018 | 0.448 | 6.850 |
| 26 | N | Echo | 0.355 | 0.046 | 0.046 | 1.710 | 0.121 | 0.149 | 0.121 | 5.180 |
| 27 | N | Bonnie | 0.028 | 0.018 | 0.411 | 0.934 | 0.093 | 0.065 | 0.093 | 6.370 |
| 28 | Y | Echo | 0.430 | 0.009 | 0.037 | 1.934 | 0.402 | 0.093 | 0.402 | 6.110 |
| 29 | Y | Bonnie | 0.046 | 0.028 | 0.168 | 0.841 | 0.261 | 0.121 | 0.261 | 6.630 |
| 30 | Y | Echo | 0.504 | 0.028 | 0.009 | 2.131 | 0.504 | 0.112 | 0.504 | 6.250 |
| 31 | N | Bonnie | 0.046 | 0.009 | 0.046 | 0.841 | 0.280 | 0.046 | 0.280 | 6.050 |
| 32 | Y | Mix | 0.392 | 0.009 | 0.084 | 0.990 | 0.168 | 0.046 | 0.168 | 7.000 |

| **Mesocosm Number** | ***Bythotrephes* Treatment** | **Stocking Lake** | ***Daphnia*** | **Small Cladocerans** | **Large Cladocerans** | **Copepodites** | **Nauplii** | **Calanoids** | **Cyclopoids** | **Total Algae** |
|---|---|---|---|---|---|---|---|---|---|---|
| 33 | Y | Bonnie | 0.018 | 0.009 | 0.028 | 1.953 | 0.168 | 0.009 | 0.168 | 6.870 |
| 34 | N | Mix | 0.289 | 0.009 | 0.084 | 1.467 | 0.196 | 0.074 | 0.196 | 6.680 |
| 35 | Y | Echo | 0.271 | 0.028 | 0.018 | 1.392 | 0.458 | 0.093 | 0.458 | 6.810 |
| 36 | N | Echo | 0.327 | 0.037 | 0.074 | 2.411 | 0.626 | 0.046 | 0.626 | 7.950 |
| 37 | N | Echo | 0.467 | 0.102 | 0.037 | 2.327 | 0.504 | 0.159 | 0.504 | 6.660 |
| 38 | Y | Echo | 0.121 | 0.046 | 0.009 | 1.280 | 0.514 | 0.037 | 0.514 | 7.650 |
| 39 | Y | Mix | 0.514 | 0.046 | 0.280 | 1.701 | 0.336 | 0.056 | 0.336 | 7.010 |
| 40 | N | Bonnie | 0.028 | 0.009 | 0.392 | 0.738 | 0.187 | 0.102 | 0.187 | 4.550 |
| **Mean ± SE** | | | 0.254±0.006 | 0.028±0.001 | 0.102±0.002 | 1.268±0.017 | 0.203±0.004 | 0.060±0.001 | 0.204±0.004 | 6.438±0.110 |
| | | Echo | 2.492±1.040 | | | *Bythotrephes* Absent | 1.954±1.026 | | | |
| | | Bonnie | 1.497±0.974 | | | *Bythotrephes* Present | 2.287±1.126 | | | |
| | | Mix | 2.191±0.861 | | | | | | | |

Table S3: Final densities (individuals per litre) in each mesocosm for *Daphnia*, small cladocerans, large cladocerans, calanoids, cyclopoids, copepodids, and nauplii prior to the application of the *Bythotrephes* treatment (N = absent, Y = present). Mean ± standard error (SE) for each taxonomic group is also provided. Densities for zooplankton taxa are provided in individuals per litre while total algal density is provided in micrograms per litre.

| **Mesocosm Number** | ***Bythotrephes* Treatment** | **Stocking Lake** | ***Daphnia*** | **Small Cladocerans** | **Large Cladocerans** | **Copepodites** | **Nauplii** | **Calanoids** | **Cyclopoids** | **Total Algae** |
|---|---|---|---|---|---|---|---|---|---|---|
| 1 | N | Mix | 0.037 | 0.028 | 0.028 | 0.373 | 1.215 | 0.009 | 1.215 | 4.780 |
| 2 | N | Echo | 0.317 | 0.074 | 0.018 | 0.420 | 0.439 | 0.009 | 0.439 | 6.750 |
| 3 | N | Bonnie | 0.159 | 1.000 | 0.018 | 0.607 | 0.626 | 0.018 | 0.626 | 6.440 |
| 4 | Y | Bonnie | 0.056 | 0.467 | 0.056 | 0.635 | 0.663 | 0.009 | 0.663 | 6.650 |
| 5 | Y | Bonnie | 0.112 | 0.934 | 0.037 | 0.570 | 0.308 | 0.009 | 0.308 | 5.850 |
| 6 | Y | Mix | 0.355 | 0.065 | 0.009 | 0.495 | 0.514 | 0.018 | 0.514 | 6.690 |
| 7 | N | Bonnie | 0.289 | 1.495 | 0.037 | 0.327 | 0.607 | 0.009 | 0.607 | 5.330 |
| 8 | Y | Echo | 0.972 | 0.233 | 0.018 | 0.383 | 0.542 | 0.037 | 0.542 | 7.100 |
| 9 | Y | Bonnie | 0.130 | 0.701 | 0.093 | 0.486 | 0.476 | 0.009 | 0.476 | 5.920 |
| 10 | N | Bonnie | 0.402 | 1.308 | 0.037 | 0.383 | 0.888 | 0.028 | 0.888 | 5.140 |
| 11 | N | Mix | 1.037 | 0.205 | 0.009 | 0.598 | 0.486 | 0.009 | 0.486 | 7.370 |
| 12 | N | Echo | 1.364 | 0.205 | 0.028 | 0.336 | 0.289 | 0.009 | 0.289 | 7.380 |
| 13 | Y | Echo | 1.140 | 0.355 | 0.018 | 0.766 | 2.869 | 0.028 | 2.869 | 6.620 |
| 14 | Y | Echo | 0.364 | 0.168 | 0.009 | 0.261 | 0.738 | 0.009 | 0.738 | 6.600 |
| 15 | N | Bonnie | 0.392 | 1.682 | 0.046 | 0.579 | 0.747 | 0.009 | 0.747 | 5.370 |
| 16 | Y | Bonnie | 0.028 | 0.224 | 0.065 | 0.542 | 0.691 | 0.028 | 0.691 | 7.020 |
| 17 | N | Bonnie | 0.177 | 0.345 | 0.028 | 0.514 | 0.719 | 0.009 | 0.719 | 5.660 |
| 18 | Y | Mix | 0.402 | 0.046 | 0.009 | 0.888 | 0.588 | 0.009 | 0.588 | 8.130 |
| 19 | N | Mix | 1.243 | 0.373 | 0.009 | 0.542 | 0.392 | 0.009 | 0.392 | 7.340 |
| 20 | N | Echo | 0.906 | 0.187 | 0.018 | 0.486 | 0.532 | 0.009 | 0.532 | 8.060 |
| 21 | Y | Bonnie | 0.028 | 0.028 | 0.056 | 0.402 | 0.831 | 0.018 | 0.831 | 8.090 |
| 22 | N | Echo | 0.588 | 0.364 | 0.018 | 0.392 | 0.719 | 0.009 | 0.719 | 9.080 |
| 23 | Y | Bonnie | 0.065 | 0.037 | 0.233 | 0.532 | 0.766 | 0.028 | 0.766 | 24.030 |
| 24 | Y | Echo | 1.430 | 0.495 | 0.037 | 0.271 | 2.822 | 0.018 | 2.822 | 7.800 |
| 25 | N | Echo | 2.280 | 0.850 | 0.084 | 0.327 | 1.149 | 0.009 | 1.149 | 7.860 |
| 26 | N | Echo | 1.327 | 0.663 | 0.018 | 0.308 | 0.953 | 0.018 | 0.953 | 7.890 |
| 27 | N | Bonnie | 0.504 | 1.289 | 0.037 | 0.869 | 0.813 | 0.056 | 0.813 | 7.200 |
| 28 | Y | Echo | 0.228 | 0.121 | 0.030 | 0.464 | 3.089 | 0.009 | 3.089 | 8.120 |
| 29 | Y | Bonnie | 0.147 | 0.323 | 0.048 | 0.712 | 0.784 | 0.009 | 0.784 | 8.110 |
| 30 | Y | Echo | 0.560 | 0.308 | 0.018 | 0.215 | 0.990 | 0.009 | 0.990 | 8.410 |
| 31 | N | Bonnie | 0.149 | 1.691 | 0.121 | 0.859 | 1.570 | 0.009 | 1.570 | 6.810 |
| 32 | Y | Mix | 1.747 | 0.159 | 0.009 | 0.317 | 0.859 | 0.009 | 0.859 | 8.420 |

| Mesocosm Number | *Bythotrephes* Treatment | Stocking Lake | *Daphnia* | Small Cladocerans | Large Cladocerans | Copepodites | Nauplii | Calanoids | Cyclopoids | Total Algae |
|---|---|---|---|---|---|---|---|---|---|---|
| 33 | Y | Bonnie | 0.074 | 0.093 | 0.009 | 0.243 | 0.065 | 0.009 | 0.065 | 10.730 |
| 34 | N | Mix | 0.944 | 0.383 | 0.009 | 0.523 | 0.850 | 0.009 | 0.850 | 8.570 |
| 35 | Y | Echo | 0.588 | 0.159 | 0.009 | 0.430 | 0.691 | 0.009 | 0.691 | 9.430 |
| 36 | N | Echo | 2.617 | 0.579 | 0.028 | 0.476 | 2.617 | 0.018 | 2.617 | 9.490 |
| 37 | N | Echo | 1.177 | 0.261 | 0.028 | 0.243 | 1.084 | 0.009 | 1.084 | 8.340 |
| 38 | Y | Echo | 0.345 | 0.261 | 0.037 | 0.411 | 1.439 | 0.009 | 1.439 | 9.000 |
| 39 | Y | Mix | 0.757 | 0.439 | 0.009 | 0.785 | 0.514 | 0.018 | 0.514 | 8.010 |
| 40 | N | Bonnie | 0.420 | 0.953 | 0.046 | 1.000 | 0.747 | 0.018 | 0.757 | 7.950 |
| **Mean ± SE** | | | 0.646 ±0.099 | 0.489 ±0.074 | 0.037 ±0.006 | 0.499 ±0.031 | 0.942 ±0.112 | 0.015 ±0.002 | 0.942 ±0.112 | 7.839 ±0.463 |

Table S4: Number of mesocosms in week 0 and week 3 with non-detections for major six major taxonomic groups and most common zooplankton species. For mesocosms with non-detections, a minimum detection density was added prior to calculation of per capita change in density.  There were no non-detections for total algae, green algae, cyanobacteria, cryptophyte, or diatom biomass in either week 0 or week 3.

| **Taxonomic groups** | **Week 0** | **Week 3** |
|---|---|---|
| *Daphnia* | 3 | None |
| Small cladocerans | 14 | None |
| Large cladocerans | 3 | 10 |
| Calanoids | 4 | 26 |
| Cyclopoids | 1 | None |
| Copepodids | None | None |
| Nauplii | None | None |
| **Most common species** | **Week 0** | **Week 3** |
| *Daphnia mendotae* | 13 | 3 |
| *Daphnia catawba* | 4 | 3 |
| *Bosmina freyi/leideri* | 21 | 5 |
| *Eubosmina tubicen* | 25 | 5 |
| *Eubosmina longispina* | 38 | 6 |
| *Skistodiaptomus oregonensis* | 40 | 10 |
| *Cyclops scutifer* | 6 | 4 |

Table S5: Estimates ± standard error (SE), degrees of freedom (df), p-values (p), and standardized estimates for all paths (represented as separate rows of response and predictor variables) assessed in piecewise structural equation model for per capita change in *Daphnia*, small cladocerans, juvenile copepods, calanoids, cyclopoids, and large cladocerans density, and total algal biomass in week 3 as response variables. Statistically significant paths ($p \leq 0.05$) are presented in bold.

| Response | Predictor | Estimate(± SE) | df | p | Standardized Estimate |
|---|---|---|---|---|---|
| ***Daphnia*** | ***Bythotrephes*** | **-1.76±0.65** | **37** | **0.0102** | **-0.11** |
| ***Daphnia*** | **Proportion of total hypolimnetic *Daphnia* in week 0** | **3.94±1.06** | **37** | **0.0006** | **0.06** |
| **Small Cladocerans** | ***Bythotrephes*** | **-1.23±0.56** | **37** | **0.0368** | **-0.01** |
| **Small Cladocerans** | **Proportion of total hypolimnetic *Daphnia* in week 0** | **2.52±1.14** | **37** | **0.0334** | **0.008** |
| Juvenile Copepods | *Bythotrephes* | -0.08±0.20 | 37 | 0.6863 | -0.05 |
| **Juvenile Copepods** | **Proportion of total hypolimnetic *Daphnia* in week 0** | **1.38±0.6541** | **37** | **0.0411** | **0.21** |
| Calanoids | *Bythotrephes* | -2.59±4.14 | 36 | 0.5362 | -2.38 |
| Calanoids | Proportion of total hypolimnetic *Daphnia* in week 0 | -21.74±32.49 | 36 | 0.5077 | -5.24 |
| Calanoids | Juvenile Copepods | 2.15±3.11 | 36 | 0.4937 | 3.41 |
| Large Cladocerans | *Bythotrephes* | 0.38±0.41 | 37 | 0.366 | 0.18 |
| Large Cladocerans | Proportion of total hypolimnetic *Daphnia* in week 0 | 1.10±1.23 | 37 | 0.375 | 0.13 |
| **Cyclopoids** | ***Bythotrephes*** | **-3.80±0.61** | **36** | **<0.0001** | **-0.11** |
| **Cyclopoids** | **Proportion of total hypolimnetic *Daphnia* in week 0** | **-35.48±6.67** | **36** | **<0.0001** | **-0.28** |
| **Cyclopoids** | **Juvenile Copepods** | **3.69±0.73** | **36** | **<0.0001** | **0.19** |
| Total Algae | *Bythotrephes* | -0.05±0.13 | 32 | 0.9626 | -0.06 |
| **Total Algae** | **Proportion of total hypolimnetic *Daphnia* in week 0** | **1.09±0.32** | **32** | **0.0017** | **0.05** |
| Total Algae | *Daphnia* | -0.04±0.01 | 32 | 0.6290 | -0.01 |
| Total Algae | Small Cladocerans | -0.002±0.001 | 32 | 0.1285 | -0.03 |
| **Total Algae** | **Large Cladocerans** | **0.18±0.04** | **32** | **<0.0001** | **0.06** |
| Total Algae | Calanoids | -0.04±0.12 | 32 | 0.7466 | -0.007 |
| **Total Algae** | **Juvenile copepods** | **-0.19±0.08** | **32** | **0.0039** | **-0.05** |

Table S6: Estimates ± standard error (SE), degrees of freedom (df), p-values (p), and standardized estimates for all paths (represented as separate rows of response and predictor variables) assessed in piecewise structural equation model with per capita change in *Daphnia*, small cladocerans, juvenile copepods, calanoids, cyclopoids, and large cladocerans density, and green algal biomass in week 3 as response variables. Statistically significant paths (p ≤ 0.05) are presented in bold.

| Response | Explanatory | Estimate(± SE) | df | p | Standardized estimate |
|---|---|---|---|---|---|
| ***Daphnia*** | ***Bythotrephes*** | **-1.76±0.65** | **37** | **0.0102** | **-0.11** |
| ***Daphnia*** | **Proportion of total hypolimnetic *Daphnia* in week 0** | **3.94±1.06** | **37** | **0.0006** | **0.06** |
| **Small Cladocerans** | ***Bythotrephes*** | **-1.22±0.56** | **37** | **0.0368** | **-0.01** |
| **Small Cladocerans** | **Proportion of total hypolimnetic *Daphnia* in week 0** | **2.52±1.14** | **37** | **0.0334** | **0.01** |
| Juvenile Copepods | *Bythotrephes* | -0.08±0.20 | 37 | 0.6863 | -0.05 |
| **Juvenile Copepods** | **Proportion of total hypolimnetic *Daphnia* in week 0** | **1.38±0.65** | **37** | **0.0411** | **0.21** |
| Calanoids | *Bythotrephes* | -2.59±4.14 | 36 | 0.5362 | -2.38 |
| Calanoids | Proportion of total hypolimnetic *Daphnia* in week 0 | -21.74±32.49 | 36 | 0.5077 | -5.24 |
| Calanoids | Juvenile Copepods | 2.15±3.11 | 36 | 0.4937 | 3.41 |
| Large Cladocerans | *Bythotrephes* | 0.38±0.41 | 37 | 0.366 | 0.18 |
| Large Cladocerans | Proportion of total hypolimnetic *Daphnia* in week 0 | 1.10±1.23 | 37 | 0.375 | 0.13 |
| **Cyclopoids** | ***Bythotrephes*** | **-3.80±0.61** | **36** | **<0.0001** | **-0.11** |
| **Cyclopoids** | **Proportion of total hypolimnetic *Daphnia* in week 0** | **-35.48±6.67** | **36** | **<0.0001** | **-0.28** |
| **Cyclopoids** | **Juvenile Copepods** | **3.69±0.72** | **36** | **<0.0001** | **0.19** |
| Green Algae | *Bythotrephes* | 0.30±0.19 | 32 | 0.13 | 0.18 |
| **Green Algae** | **Proportion of total hypolimnetic *Daphnia* in week 0** | **2.92±0.37** | **32** | **<0.0001** | **0.44** |
| Green Algae | *Daphnia* | -0.02±0.01 | 32 | 0.87832 | -0.02 |
| Green Algae | Small Cladocerans | 0.0009±0.001 | 32 | 0.562 | 0.04 |
| **Green Algae** | **Large Cladocerans** | **0.36±0.04** | **32** | **<0.0001** | **0.43** |
| Green Algae | Calanoids | 0.25±0.15 | 32 | 0.112 | 0.16 |
| **Green Algae** | **Juvenile copepods** | **-0.14±0.07** | **32** | **0.0436** | **-0.14** |

Table S7: Estimates ± standard error (SE), degrees of freedom (df), p-values (p), and standardized estimates for all paths (represented as separate rows of response and predictor variables) assessed in piecewise structural equation model with per capita change in *Daphnia*, small cladocerans, juvenile copepods, calanoids, cyclopoids, and large cladocerans density, and diatom biomass in week 3 as response variables. Statistically significant paths ($p \leq 0.05$) are presented in bold.

| Response | Explanatory | Estimate(± SE) | df | p | Standardized estimate |
|---|---|---|---|---|---|
| ***Daphnia*** | ***Bythotrephes*** | **-1.76±0.65** | **37** | **0.0102** | **-0.11** |
| ***Daphnia*** | **Proportion of total hypolimnetic *Daphnia* in week 0** | **3.94±1.06** | **37** | **0.0006** | **0.06** |
| **Small Cladocerans** | ***Bythotrephes*** | **-1.22±0.56** | **37** | **0.0368** | **-0.01** |
| **Small Cladocerans** | **Proportion of total hypolimnetic *Daphnia* in week 0** | **2.52±1.14** | **37** | **0.0334** | **0.01** |
| Juvenile Copepods | *Bythotrephes* | -0.08±0.20 | 37 | 0.6863 | -0.05 |
| **Juvenile Copepods** | **Proportion of total hypolimnetic *Daphnia* in week 0** | **1.38±0.65** | **37** | **0.0411** | **0.21** |
| Calanoids | *Bythotrephes* | -2.59±4.14 | 36 | 0.5362 | -2.38 |
| Calanoids | Proportion of total hypolimnetic *Daphnia* in week 0 | -21.74±32.49 | 36 | 0.5077 | -5.24 |
| Calanoids | Juvenile Copepods | 2.15±3.11 | 36 | 0.4937 | 3.41 |
| Large Cladocerans | *Bythotrephes* | 0.38±0.41 | 37 | 0.366 | 0.18 |
| Large Cladocerans | Proportion of total hypolimnetic *Daphnia* in week 0 | 1.10±1.23 | 37 | 0.375 | 0.13 |
| **Cyclopoids** | ***Bythotrephes*** | **-3.80±0.61** | **36** | **<0.0001** | **-0.11** |
| **Cyclopoids** | **Proportion of total hypolimnetic *Daphnia* in week 0** | **-35.48±6.67** | **36** | **<0.0001** | **-0.28** |
| **Cyclopoids** | **Juvenile Copepods** | **3.69±0.72** | **36** | **<0.0001** | **0.19** |
| **Diatoms** | ***Bythotrephes*** | **-1.05±0.50** | **32** | **0.04** | **-0.44** |
| **Diatoms** | **Proportion of total hypolimnetic *Daphnia* in week 0** | **1.65±1.89** | **32** | **<0.0001** | **1.48** |
| Diatoms | *Daphnia* | -0.06±0.04 | 32 | 0.186 | -0.41 |
| **Diatoms** | **Small Cladocerans** | **-0.02±0.005** | **32** | **0.003** | **-0.62** |
| **Diatoms** | **Large Cladocerans** | **0.73±0.08** | **32** | **<0.0001** | **0.65** |
| **Diatoms** | **Calanoids** | **-3.88±0.76** | **32** | **<0.0001** | **-1.75** |
| **Diatoms** | **Juvenile copepods** | **-0.39±0.15** | **32** | **0.0152** | **-0.28** |

Table S8: Estimates ± standard error (SE), degrees of freedom (df), p-values (p), and standardized estimates for all paths (represented as separate rows of response and predictor variables) assessed in piecewise structural equation model with per capita change in *Daphnia*, small cladocerans, juvenile copepods, calanoids, cyclopoids, and large cladocerans density, and cyanobacteria biomass in week 3 as response variables. Statistically significant paths (p ≤ 0.05) are presented in bold.

| Response | Explanatory | Estimate(± SE) | df | p | Standardized estimate |
|---|---|---|---|---|---|
| ***Daphnia*** | ***Bythotrephes*** | **-1.76±0.65** | **37** | **0.0102** | **-0.11** |
| ***Daphnia*** | **Proportion of total hypolimnetic *Daphnia* in week 0** | **3.94±1.06** | **37** | **0.0006** | **0.06** |
| **Small Cladocerans** | ***Bythotrephes*** | **-1.22±0.56** | **37** | **0.0368** | **-0.01** |
| **Small Cladocerans** | **Proportion of total hypolimnetic *Daphnia* in week 0** | **2.52±1.14** | **37** | **0.0334** | **0.01** |
| Juvenile Copepods | *Bythotrephes* | -0.08±0.20 | 37 | 0.6863 | -0.05 |
| **Juvenile Copepods** | **Proportion of total hypolimnetic *Daphnia* in week 0** | **1.38±0.65** | **37** | **0.0411** | **0.21** |
| Calanoids | *Bythotrephes* | -2.59±4.14 | 36 | 0.5362 | -2.38 |
| Calanoids | Proportion of total hypolimnetic *Daphnia* in week 0 | -21.74±32.49 | 36 | 0.5077 | -5.24 |
| Calanoids | Juvenile Copepods | 2.15±3.11 | 36 | 0.4937 | 3.41 |
| Large Cladocerans | *Bythotrephes* | 0.38±0.41 | 37 | 0.366 | 0.18 |
| Large Cladocerans | Proportion of total hypolimnetic *Daphnia* in week 0 | 1.10±1.23 | 37 | 0.375 | 0.13 |
| **Cyclopoids** | ***Bythotrephes*** | **-3.80±0.61** | **36** | **<0.0001** | **-0.11** |
| **Cyclopoids** | **Proportion of total hypolimnetic *Daphnia* in week 0** | **-35.48±6.67** | **36** | **<0.0001** | **-0.28** |
| **Cyclopoids** | **Juvenile Copepods** | **3.69±0.72** | **36** | **<0.0001** | **0.19** |
| Cyanobacteria | *Bythotrephes* | 0.001±0.081 | 32 | 0.963 | 0.0022 |
| Cyanobacteria | Proportion of total hypolimnetic *Daphnia* in week 0 | -0.258±0.325 | 32 | 0.433 | -0.023 |
| Cyanobacteria | *Daphnia* | 0.001±0.007 | 32 | 0.899 | 0.004 |
| Cyanobacteria | Small Cladocerans | -0.001±0.001 | 32 | 0.243 | -0.042 |
| Cyanobacteria | Large Cladocerans | 0.046±0.037 | 32 | 0.216 | 0.033 |
| Cyanobacteria | Calanoids | -0.071±0.084 | 32 | 0.407 | -0.026 |
| Cyanobacteria | Juvenile copepods | -0.122±0.061 | 32 | 0.055 | -0.072 |

Table S9: Estimates ± standard error (SE), degrees of freedom (df), p-values (p), and standardized estimates for all paths (represented as separate rows of response and predictor variables) assessed in piecewise structural equation model with per capita change in *Daphnia*, small cladocerans, juvenile copepods, calanoids, cyclopoids, and large cladocerans density, and cryptophyte biomass in week 3 as response variables. Statistically significant paths (p ≤ 0.05) are presented in bold.

| Response | Explanatory | Estimate(± SE) | df | p | Standardized estimate |
|---|---|---|---|---|---|
| ***Daphnia*** | ***Bythotrephes*** | **-1.76±0.65** | **37** | **0.0102** | **-0.11** |
| ***Daphnia*** | **Proportion of total hypolimnetic *Daphnia* in week 0** | **3.94±1.06** | **37** | **0.0006** | **0.06** |
| **Small Cladocerans** | ***Bythotrephes*** | **-1.22±0.56** | **37** | **0.0368** | **-0.01** |
| **Small Cladocerans** | **Proportion of total hypolimnetic *Daphnia* in week 0** | **2.52±1.14** | **37** | **0.0334** | **0.01** |
| Juvenile Copepods | *Bythotrephes* | -0.08±0.20 | 37 | 0.6863 | -0.05 |
| **Juvenile Copepods** | **Proportion of total hypolimnetic *Daphnia* in week 0** | **1.38±0.65** | **37** | **0.0411** | **0.21** |
| Calanoids | *Bythotrephes* | -2.59±4.14 | 36 | 0.5362 | -2.38 |
| Calanoids | Proportion of total hypolimnetic *Daphnia* in week 0 | -21.74±32.49 | 36 | 0.5077 | -5.24 |
| Calanoids | Juvenile Copepods | 2.15±3.11 | 36 | 0.4937 | 3.41 |
| Large Cladocerans | *Bythotrephes* | 0.38±0.41 | 37 | 0.366 | 0.18 |
| Large Cladocerans | Proportion of total hypolimnetic *Daphnia* in week 0 | 1.10±1.23 | 37 | 0.375 | 0.13 |
| **Cyclopoids** | ***Bythotrephes*** | **-3.80±0.61** | **36** | **<0.0001** | **-0.11** |
| **Cyclopoids** | **Proportion of total hypolimnetic *Daphnia* in week 0** | **-35.48±6.67** | **36** | **<0.0001** | **-0.28** |
| **Cyclopoids** | **Juvenile Copepods** | **3.69±0.72** | **36** | **<0.0001** | **0.19** |
| Cryptophytes | *Bythotrephes* | 0.210±0.278 | 32 | 0.456 | 0.283 |
| Cryptophytes | Proportion of total hypolimnetic *Daphnia* in week 0 | 0.509±0.890 | 32 | 0.571 | -0.180 |
| Cryptophytes | *Daphnia* | 0.106±0.016 | 32 | 0.298 | 0.358 |
| Cryptophytes | Small Cladocerans | 0±0.003 | 32 | 0.993 | -0.042 |
| **Cryptophytes** | **Large Cladocerans** | **0.251±0.078** | **32** | **0.003** | **0.723** |
| Cryptophytes | Calanoids | -0.311±0.423 | 32 | 0.468 | -0.556 |
| Cryptophytes | Juvenile copepods | -0.384±0.205 | 32 | 0.070 | -0.893 |

Table S10: Estimates ± standard error(SE), degrees of freedom (df), p-values (p), and standardized estimates for all paths (represented as separate rows of response and predictor variables) assessed in piecewise structural equation model with per capita change in most common *Daphnia*, small cladocerans, calanoid, and cyclopoid species as response variables. Statistically significant paths (p ≤ 0.05) are presented in bold.

| Response | Explanatory | Estimate (± SE) | df | p | Standardized estimate |
|---|---|---|---|---|---|
| ***Daphnia mendotae*** | ***Bythotrephes*** | **-0.97±0.46** | **37** | **0.0429** | **-0.09** |
| *Daphnia mendotae* | Proportion of total hypolimnetic *Daphnia* in week 0 | 1.58±1.09 | 37 | 0.1554 | 0.04 |
| *Daphnia catawba* | *Bythotrephes* | -0.61±0.46 | 37 | 0.1907 | -0.10 |
| *Daphnia catawba* | Proportion of total hypolimnetic *Daphnia* in week 0 | -0.70±1.64 | 37 | 0.6715 | -0.03 |
| *Bosmina freyi/leideri* | *Bythotrephes* | -1.44±0.76 | 37 | 0.0678 | -0.02 |
| ***Bosmina freyi/leideri*** | **Proportion of total hypolimnetic *Daphnia* in week 0** | **2.72±1.33** | **37** | **0.0487** | **0.01** |
| *Eubosmina tubicen* | *Bythotrephes* | -0.73±0.50 | 37 | 0.1556 | -0.03 |
| *Eubosmina tubicen* | Proportion of total hypolimnetic *Daphnia* in week 0 | -0.65±1.66 | 37 | 0.6973 | -0.01 |
| ***Eubosmina longispina*** | ***Bythotrephes*** | **-0.49±0.24** | **37** | **0.0511** | **-0.03** |
| ***Eubosmina longispina*** | **Proportion of total hypolimnetic *Daphnia* in week 0** | **-3.19±1.39** | **37** | **0.0278** | **-0.05** |
| *Skistodiaptomus oregonensis* | *Bythotrephes* | 0.51±0.37 | 37 | 0.1744 | 0.03 |
| *Skistodiaptomus oregonensis* | Proportion of total hypolimnetic *Daphnia* in week 0 | -8.03±4.40 | 37 | 0.0761 | -0.11 |
| ***Cyclops scutifer*** | ***Bythotrephes*** | **-1.27±0.62** | **37** | **0.0458** | **-0.18** |
| ***Cyclops scutifer*** | **Proportion of total hypolimnetic *Daphnia* in week 0** | **3.55±1.35** | **37** | **0.0123** | **0.13** |

Table S11: Estimates ± standard error(SE), degrees of freedom (df), p-values (p), and standardized estimates for all paths (represented as separate rows of response and predictor variables) assessed in piecewise structural equation model fit with proportion of hypolimnetic *Daphnia mendotae* in Week 0 as a predictor variable and per capita change in *Daphnia*, small cladocerans, large cladocerans, calanoids, cyclopoids, juvenile copepods, and total algae density as response variables. Statistically significant paths ($p \leq 0.05$) are presented in bold.

| Response | Predictor | Estimate(±SE) | df | p | Standardized Estimate |
|---|---|---|---|---|---|
| ***Daphnia*** | ***Bythotrephes*** | **-3.51±1.68** | **37** | **0.0435** | **-0.22** |
| ***Daphnia*** | **Proportion of hypolimnetic *Daphnia mendotae* in week 0** | **3.86±1.33** | **37** | **0.0061** | **0.09** |
| **Small Cladocerans** | ***Bythotrephes*** | **-1.22±0.61** | **37** | **0.0522** | **-0.01** |
| Small Cladocerans | Proportion of hypolimnetic *Daphnia mendotae* in week 0 | 1.45±1.01 | 37 | 0.1584 | 0.01 |
| Juvenile Copepods | *Bythotrephes* | -0.03±0.22 | 37 | 0.8827 | -0.02 |
| Juvenile Copepods | Proportion of hypolimnetic *Daphnia mendotae* in week 0 | 0.63±0.48 | 37 | 0.2036 | 0.13 |
| Calanoids | *Bythotrephes* | -0.13±0.37 | 36 | 0.7352 | -0.12 |
| Calanoids | Proportion of hypolimnetic *Daphnia mendotae* in week 0 | -0.04±1.21 | 36 | 0.9736 | -0.01 |
| Calanoids | Juvenile Copepods | 0.26±0.17 | 36 | 0.1348 | 0.41 |
| Large Cladocerns | *Bythotrephes* | 0.46±0.44 | 37 | 0.3017 | 0.21 |
| Large Cladocerns | Proportion of hypolimnetic *Daphnia mendotae* in week 0 | 0.06±0.96 | 37 | 0.9474 | 0.01 |
| Cyclopoids | *Bythotrephes* | -0.24±0.39 | 36 | 0.5423 | -0.01 |
| Cyclopoids | Proportion of hypolimnetic *Daphnia mendotae* in week 0 | -1.18±1.71 | 36 | 0.4968 | -0.01 |
| Cyclopoids | Juvenile Copepods | 0.57±0.17 | 36 | 0.0024 | 0.03 |
| **Total Algae** | ***Bythotrephes*** | **-0.03±0.14** | **32** | **0.8426** | **-0.03** |
| Total Algae | Proportion of hypolimnetic *Daphnia mendotae* in week 0 | 0.53±0.27 | 32 | 0.0625 | 0.20 |
| Total Algae | *Daphnia* | -0.001±0.01 | 32 | 0.8926 | -0.02 |
| Total Algae | Small Cladocerans | -0.001±0.002 | 32 | 0.4533 | -0.12 |
| **Total Algae** | **Large Cladocerans** | **0.19±0.05** | **32** | **0.0004** | **0.42** |
| Total Algae | Calanoids | 0.10±0.11 | 32 | 0.3964 | 0.11 |
| **Total Algae** | **Juvenile Copepods** | **-0.20±0.09** | **32** | **0.026** | **-0.36** |

Table S12: Estimates ±SE, degrees of freedom (df), p-values (p), and standardized estimates for all paths (represented as separate rows of response and predictor variables) assessed in piecewise structural equation model fit with proportion of hypolimnetic *Daphnia catawba* in Week 0 as a predictor variable and per capita change in *Daphnia*, small cladocerans, large cladocerans, calanoids, cyclopoids, juvenile copepods, and total algae density as response variables. Statistically significant paths (p ≤ 0.05) are presented in bold.

| Response | Predictor | Estimate (±SE) | df | p | Standardized Estimate |
|---|---|---|---|---|---|
| ***Daphnia*** | ***Bythotrephes*** | **-2.41±0.89** | **37** | **0.0101** | **-0.1506** |
| ***Daphnia*** | **Proportion of hypolimnetic *Daphnia catawba* in week 0** | **3.26±0.97** | **37** | **0.0018** | **0.0718** |
| Small Cladocerans | *Bythotrephes* | -0.98±0.55 | 37 | 0.0838 | -0.0113 |
| Small Cladocerans | Proportion of hypolimnetic *Daphnia catawba* in week 0 | 0.78±1.04 | 37 | 0.46 | 0.0032 |
| Juvenile Copepods | *Bythotrephes* | -0.01±0.21 | 37 | 0.95 | -0.0079 |
| Juvenile Copepods | Proportion of hypolimnetic *Daphnia catawba* in week 0 | 0.57±0.50 | 37 | 0.2628 | 0.1176 |
| Calanoids | *Bythotrephes* | -0.33±0.39 | 36 | 0.39 | -0.3093 |
| Calanoids | Proportion of hypolimnetic *Daphnia catawba* in week 0 | -3.11±2.43 | 36 | 0.2081 | -1.0134 |
| **Calanoids** | **Juvenile copepods** | **0.52±0.25** | **36** | **0.0404** | **0.8317** |
| Large Cladocerans | *Bythotrephes* | 0.46±0.43 | 37 | 0.302 | 0.2144 |
| Large Cladocerans | Proportion of hypolimnetic *Daphnia catawba* in week 0 | 0.23±0.96 | 37 | 0.8071 | 0.039 |
| **Cyclopoids** | ***Bythotrephes*** | **-1.39±0.51** | **36** | **0.0097** | **-0.0415** |
| **Cyclopoids** | **Proportion of hypolimnetic *Daphnia catawba* in week 0** | **-13.99±4.38** | **36** | **0.0029** | **-0.1474** |
| Cyclopoids | Juvenile copepods | 1.62±0.47 | 36 | 0.0015 | 0.0833 |
| Total Algae | *Bythotrephes* | -0.02±0.14 | 32 | 0.884 | -0.0208 |
| **Total Algae** | **Proportion of hypolimnetic *Daphnia catawba* in week 0** | **0.58±0.27** | **32** | **0.0444** | **0.2085** |
| Total Algae | *Daphnia* | -0.005±0.01 | 32 | 0.6432 | -0.0857 |
| Total Algae | Small Cladocerans | -0.001±0.002 | 32 | 0.5906 | -0.0825 |
| **Total Algae** | **Large Cladocerans** | **0.18±0.05** | **32** | **0.0007** | **0.3965** |
| Total Algae | Calanoids | 0.05±0.12 | 32 | 0.6952 | 0.0539 |
| **Total Algae** | **Juvenile copepods** | **-0.17±0.08** | **32** | **0.0339** | **-0.3089** |

Table S13: Estimates ± standard error (SE), degrees of freedom (df), p-values (p) for generalized linear models or robust regression fitted with a gamma distribution and a log-link function assessing the interactive effect of *Bythotrephes* presence and proportion of total hypolimnetic *Daphnia* in week 0 on the per capita change in density of zooplankton taxonomic groups and species, and biomass of major algal groups with significant paths for one or both of these variables in piecewise structural equation models. P-values were determined from log-likelihood ratio tests with a Chi-squared distribution for GLMs and Wald test for a robust regression.

| Response Variable | df | Estimate (±SE) | p | Analysis |
|---|---|---|---|---|
| *Daphnia* | 36 | -3.63±3.61 | 0.02 | Gamma GLM with log link |
| *Daphnia mendotae* | 36 | -3.30±2.24 | 0.60 | Gamma GLM with log link |
| Small Cladocerans | 36 | -6.48±2.67 | 0.02 | Gamma GLM with log link |
| *Bosmina freyi/leideri* | 36 | -4.66±3.48 | 0.50 | Gamma GLM with log link |
| *Eubosmina longispina* | 36 | -1.40±1.78 | 0.30 | Gamma GLM with log link |
| Cyclopoids | 36 | -0.67±3.97 | 0.82 | Gamma GLM with log link |
| *C. scutifer* | 36 | -8.41±3.27 | 0.02 | Gamma GLM with log link |
| Total Algae | 37 | 1.47±0.39 | 0.0002 | Gamma Robust Regression with log link |
| Green Algae | 37 | 0.81±1.5 | 0.60 | Gamma GLM with log link |
| Diatoms | 37 | 2.00±3.12 | 0.52 | Gamma Robust Regression with log link |

Table S14: Summary of results from Gamma distributed GLMMs assessing the differences in the proportion hypo- (Hypo) *Daphnia* between invaded and uninvaded mesocosms (Byth; N or Y) across experimental weeks 0 and 3 (Week; 0 or 3). FDR refers to the false discovery rate adjusted p-values with statistical significance denoted in brackets. Generalized linear hypothesis tests performed (GLHT) for relevant comparisons when significant interactions between predictor variables were detected. None refers to no significant predictors detected, with no relevant p values reported (NA). Degrees of freedom (df) are provided in brackets, next to p-values.

| **Taxa/species** | **Response variable** | **Significant Predictor(s)** | **p-value (df)** | **Non-significant predictors** | **GLHT comparisons** | **p-value** |
|---|---|---|---|---|---|---|
| Total *Daphnia* | Hypo | Byth x Week | 0.03 (76) | None | N0-N3<br>Y0-Y3<br>N3-Y3 | 0.07<br>0.09<br>0.07 |
| *D. catawba* | Hypo | None | NA | Byth x Week, Byth, Week | None | NA |
| *D. mendotae* | Hypo | Byth x Week | 0.03 (76) | None | N0-N3<br>Y0-Y3<br>N3-Y3 | 0.018<br>0.698<br>0.09 |

**Figures**

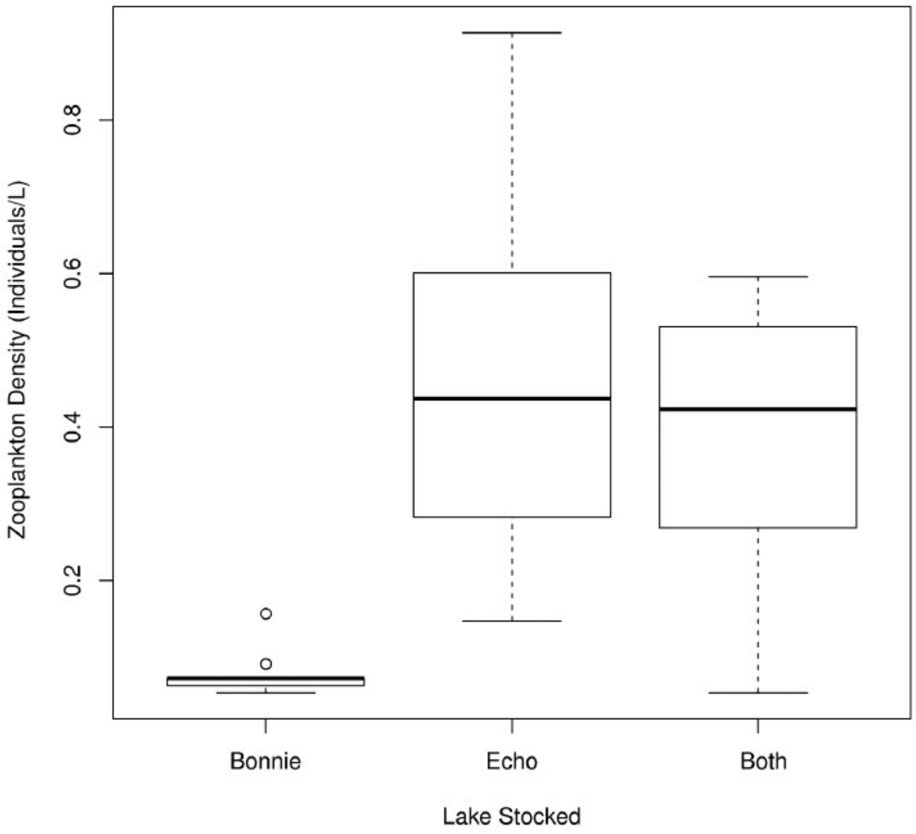


Figure S1: Zooplankton density (individuals/L) in mesocosms stocked from Echo, Bonnie or Both lakes prior to the start of the experiment.

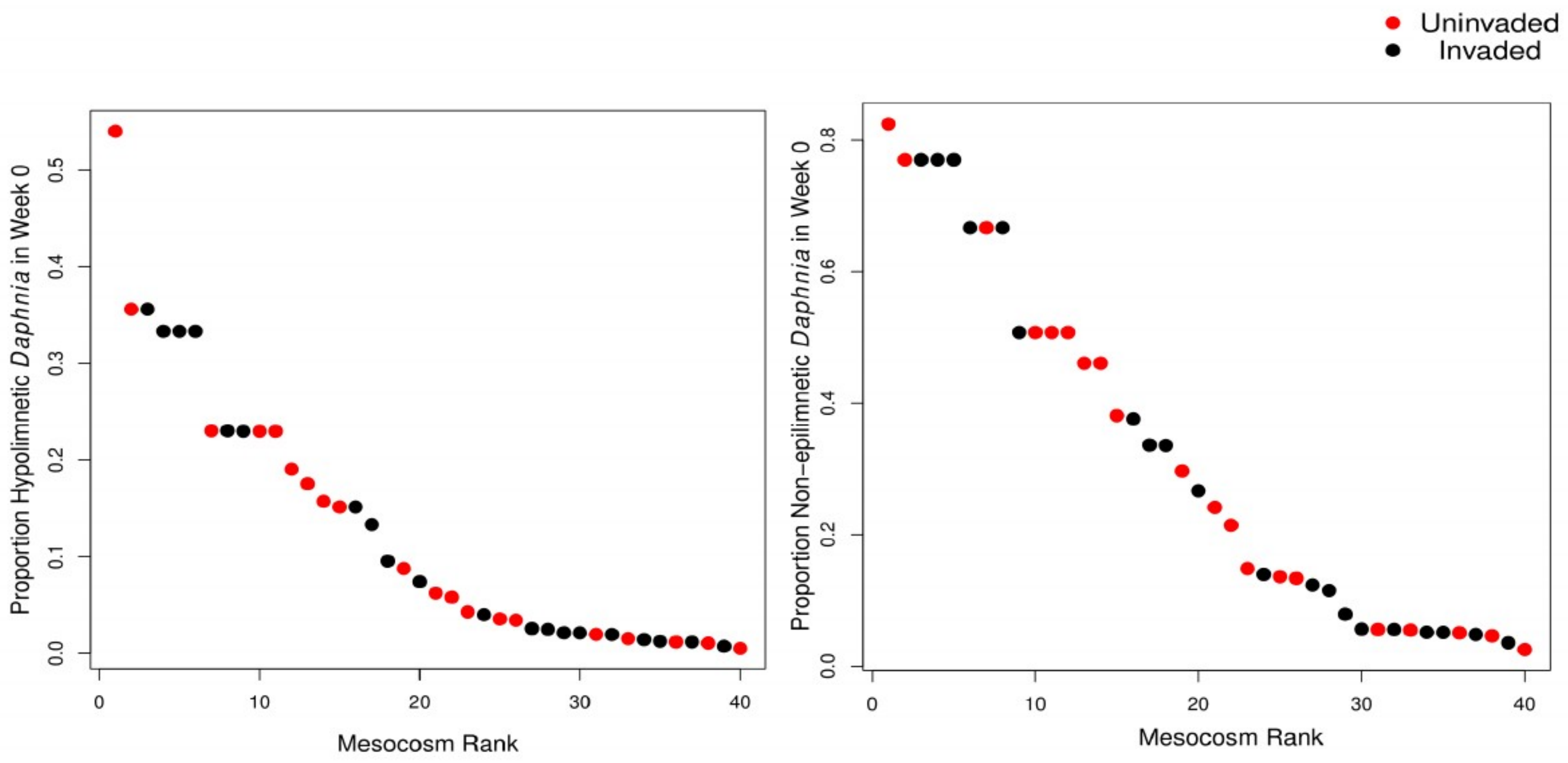


Figure S2: The proportion of hypo- and non-epilimnetic *Daphnia* across all forty mesocosms prior to the start of the experiment (week 0). Red dots indicate mesocosms invaded by *Bythotrephes* in week 1, while black dots indicate mesocosms which remained uninvaded.

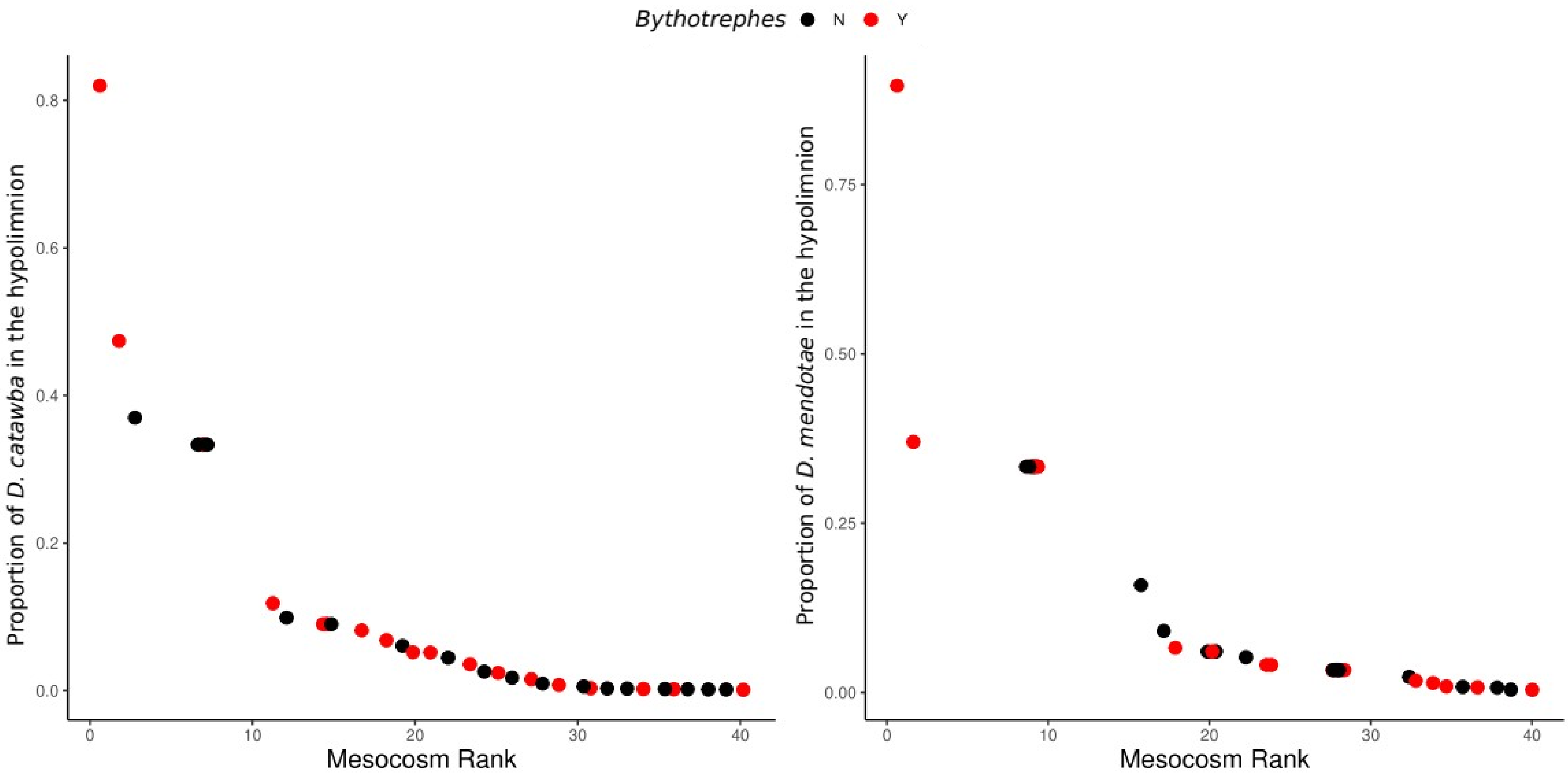


Figure S3: The proportion of hypolimnetic *D. catawba* and *D. mendotae* across all forty mesocosms prior to the start of the experiment (week 0). Red dots indicate mesocosms invaded by *Bythotrephes* in week 1, while black dots indicate mesocosms which remained uninvaded.

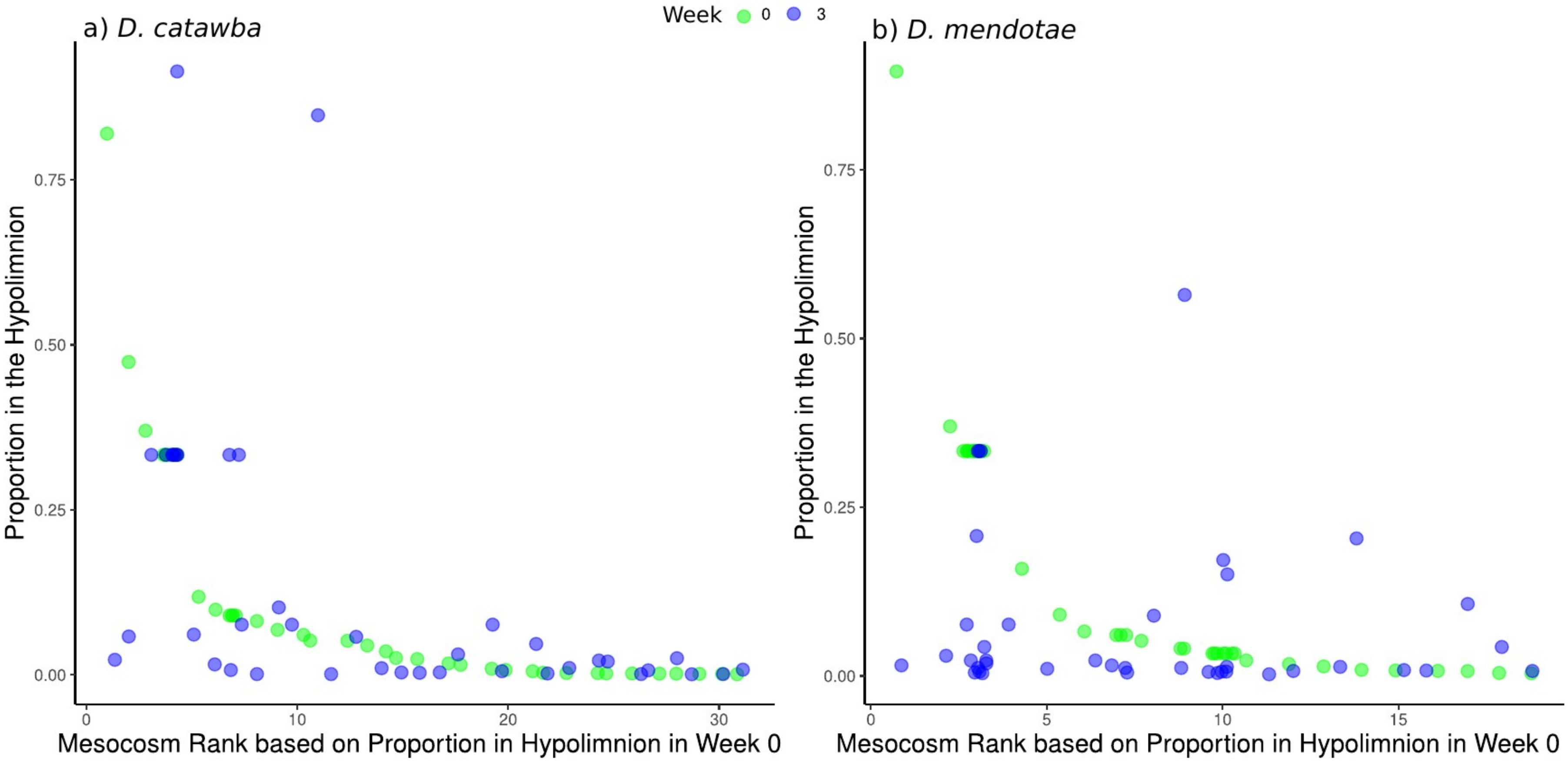


Figure S4: The proportion of hypolimnetic *D. catawba* and *D. mendotae* across all forty mesocosms in Week 0 (green) and Week 3 (blue).

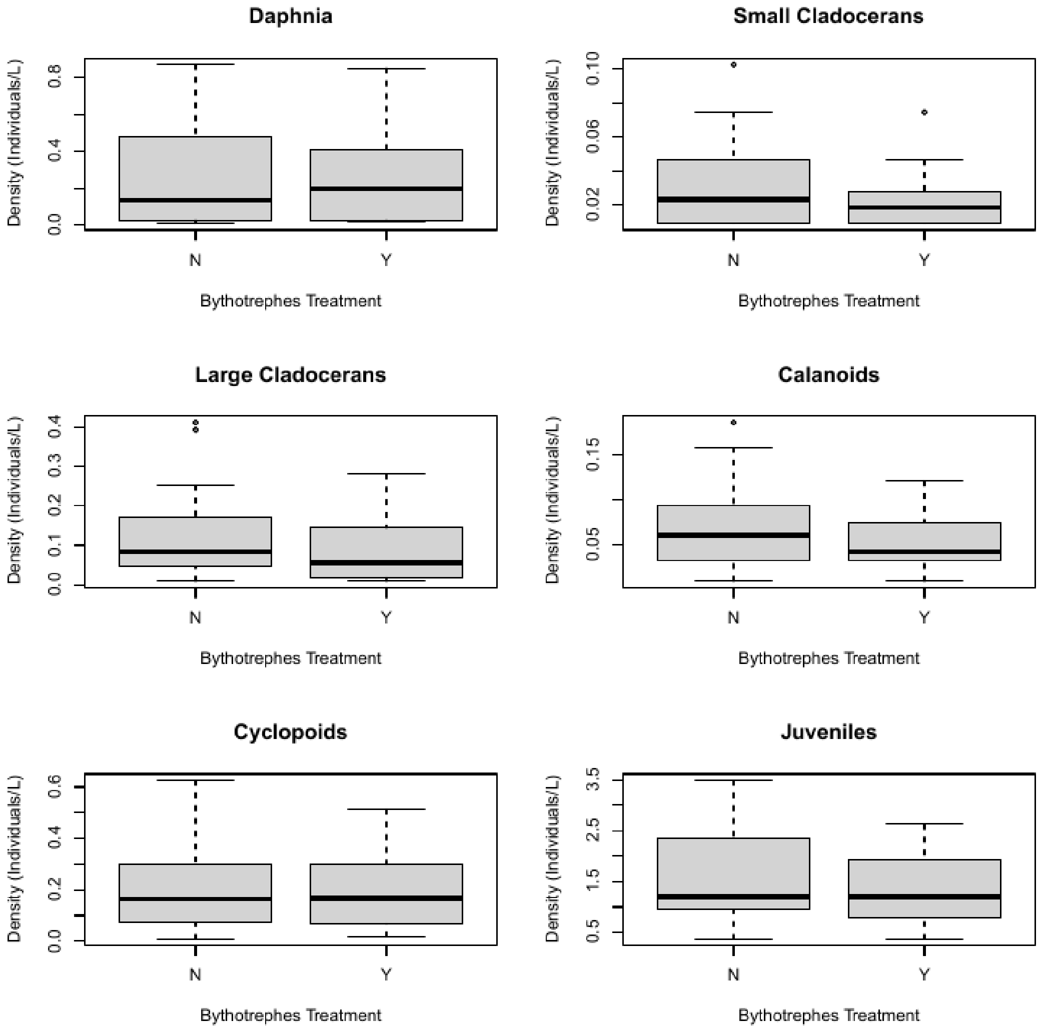


Figure S5: Starting densities (week 0) of total *Daphnia*, small cladoceran, large cladocerans, calanoids, cyclopoids, and juveniles in mesocosms assigned *Bythotrephes* invaded (Y) and uninvaded (N) treatments (linear model with *Bythotrephes* as a predictor variable, *Daphnia*; p = 0.781, small cladoceran; p = 0.229, carge cladocerans; p = 0.186, calanoids; p = 0.224, cyclopoids; p = 0.91, and juveniles; p = 0.448). Week 0 densities were measured prior to the stocking of *Bythotrephes* in invaded treatments.

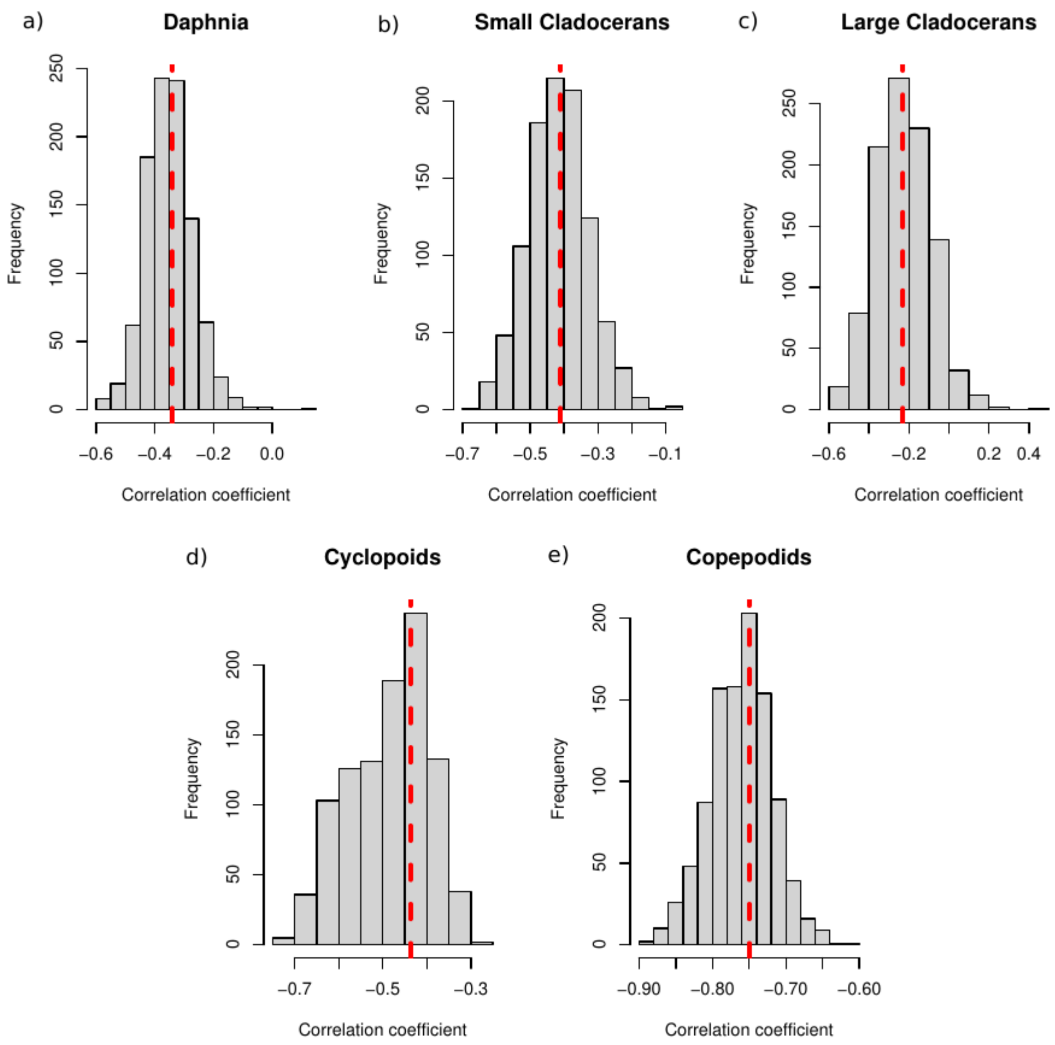


Figure S6: Sampling distributions of bootstrapped correlation coefficients between per capita change in density for a) *Daphnia*, b) small cladocerans, c) large cladocerans, d) cyclopoids, and copepodids. Dashed line represents the value of the coefficient obtained from correlation analysis between taxa per capita change in density and total zooplankton density (individuals/L) in week 0.

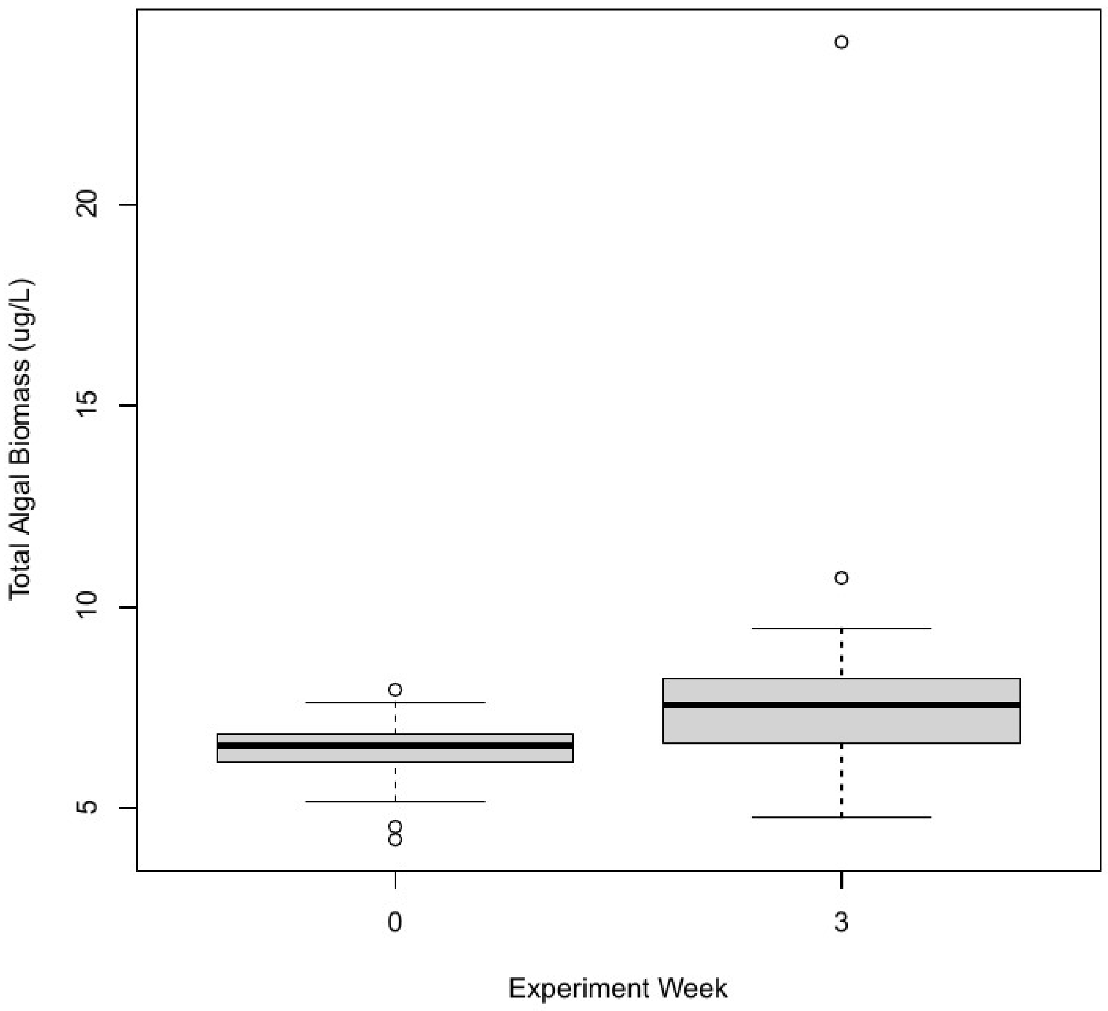


Figure S7: Differences in total algal biomass (µg/L) between the experiment week 0 and week 3 across all mesocosms. Total algal biomass was significantly greater in week 3 as compared to week 0 (two sample t-test, df = 43.4, p = 0.006).
